# Antecedent effect models as an exploratory tool to link climate drivers to herbaceous perennial population dynamics data

**DOI:** 10.1101/2022.03.11.484031

**Authors:** Aldo Compagnoni, Dylan Childs, Tiffany M. Knight, Roberto Salguero- Gómez

**Affiliations:** Martin Luther University Halle-Wittenberg, Am Kirchtor 1, 06108, Halle (Saale), Germany; Department of Community Ecology, Helmholtz Centre for Environmental Research–UFZ, 06120 Halle (Saale), Germany; Deutscher Platz 5e, Leipzig, 04103, Germany; German Centre for Integrative Biodiversity Research (iDiv) Halle-Jena-Leipzig, Puschstrasse 4, Leipzig 04103, Germany; Deutscher Platz 5e, Leipzig, 04103, Germany; Department of Animal and Plant Sciences, University of Sheffield. Western Bank, Sheffield S10 2TN, UK; Deutscher Platz 5e, Leipzig, 04103, Germany; Department of Biology, University of Oxford. South Parks Road, Oxford, OX1 3SZ, United Kingdom; Deutscher Platz 5e, Leipzig, 04103, Germany

**Keywords:** Bayesian moving window, environmental drivers, demography, lag, lagged climate effect, delay, COMPADRE Plant Matrix Database, cross-validation, forecasting, population growth rate, regularization, vital rate

## Abstract

Understanding mechanisms and predicting natural population responses to climate is a key goal of Ecology. However, studies explicitly linking climate to population dynamics remain limited. Antecedent effect models are a set of statistical tools that capitalize on the evidence provided by climate and population data to select time windows correlated with a response (*e.g.*, survival, reproduction). Thus, these models can serve as both a predictive and exploratory tool. We compare the predictive performance of antecedent effect models against simpler models, and showcase their exploratory analysis potential by selecting a case study with high predictive power. We fit three antecedent effect models: (1) weighted mean models (WMM), which weigh the importance of monthly anomalies based on a Gaussian curve, (2) stochastic antecedent models (SAM), which weigh the importance of monthly anomalies using a Dirichlet process, and (3) regularized regressions using the Finnish Horseshoe prior (FHM), which estimate a separate effect size for each monthly anomaly. We compare these approaches to a linear model using a yearly climatic predictor and a null model with no predictors. We use demographic data from 77 natural populations of 34 plant species ranging between seven and 36 years of length. We then fit models to the asymptotic population growth rate (λ) and its underlying vital rates: survival, development, and reproduction. We find that models including climate do not consistently outperform null models. We hypothesize that the effect of yearly climate is too complex, weak, and confounded by other factors to be easily predicted using monthly precipitation and temperature data. On the other hand, in our case study, antecedent effect models show biologically sensible correlations between two precipitation anomalies and multiple vital rates. We conclude that, in temporal datasets with limited sample sizes, antecedent effect models are better suited as exploratory tools for hypothesis generation.

**Open Research statement:** Data and code to reproduce the analyses are available on zenodo at https://dx.doi.org/10.5281/zenodo.7839199.

## Introduction

Understanding and predicting population dynamics is a key objective of Ecology (Sutherland et al. 2013). In population ecology, the last decades have witnessed the development and application of many statistical methods aimed precisely at this objective (Caswell 2001, Ellner et al. 2016). Increasingly, emphasis is being devoted to understanding how climatic drivers such as temperature and precipitation affect vital rates like survival, development (*i.e.,* changes in stage along the life cycle), and reproduction (Ehrlén et al. 2016; Teller et al. 2016). The ultimate goal of this ambitious research agenda is to achieve a mechanistic understanding of what makes species vulnerable to climate change worldwide (Sutherland et al. 2013). However, progress in this understanding is underpinned by synthetic studies that explicitly link vital rates to climatic drivers using standardized methods (Ehrlén et al. 2016, Compagnoni et al. 2021).

To understand and predict the effects of climate drivers on populations, ecologists need to estimate the predictability of different response variables. While ecologists are ultimately interested in linking population performance metrics (*e.g.*, population growth rate) to climatic drivers, this link often has poor predictive ability (Knape and de Valpine 2011, Tredennick et al. 2016). The low predictive performance might occur because *a priori* knowledge of the system is often poor, and/or because the number of potential covariates is orders of magnitude larger than the number of data points available in population studies (Tredennick et al. 2021). In addition, poor predictive performance might reflect how variation in population growth rates might be buffered against environmental stochasticity, in that responses of single vital rates might not fully translate into population growth rate responses (Hilde et al. 2020). Thus, we might have a better understanding and ability to predict the effects of climate on vital rates (*e.g.*, Clark et al., 2016, 2021; Schulze et al., 2019). Alternatively, the low predictive ability of our models might simply reflect how the approaches to link populations dynamics to climate implemented so far are ineffective. Currently, we lack a key piece of the puzzle: the relative ability of existing approaches to predict vital rates and population growth rate, and how this predictability varies across species. Establishing this predictive ability in synthesis study requires addressing the technical hurdles associated with linking climatic drivers to population dynamics.

Studies that explicitly link population responses to climate drivers typically identify climatic predictors *a priori*. Most population studies are short-term (Salguero-Gómez et al. 2015, 2016, Römer et al. 2021, Levin et al. 2022), thus preventing investigators from effectively using exploratory or model selection approaches. As a result, ecologist have historically considered seasonal or, more frequently, annual climatic effects (reviewed in Evers et al., 2021). This choice has practical reasons: first, monthly or annual climate data are widely available (Daly et al. 1994, Karger and Zimmermann 2018). Second, annual anomalies are a logical predictor of annual demographic variation, and seasonal anomalies can be justified by physiological or behavioural considerations (White et al. 1999, Catchpole et al. 2000). However, no consensus on these *a priori* choices exists. For example, expectations based on physiology, such as the importance of the growing season in plants, or the importance of exceeding minimum thermal limits in insects (Angilletta, 2009), are often met with opposing evidence (Kreyling 2010, Czachura and Miller 2020), or evidence of lagged effects from previous years (Hacket Pain et al. 2018, Tenhumberg et al. 2018, Evers et al. 2021). At worst, models that link population responses to monthly or annual climatic variables could be an excessive oversimplification of the processes that drive plant physiology. The effect of climate on plants unfolds via the dynamic, nonlinear interaction of temperature, soil moisture, relative humidity, and radiative fluxes at sub-hourly time scales (Lambers et al. 2008). While these factors correlate with annual or seasonal temperature and precipitation, such correlation might be too weak to make annual or seasonal climate effective predictors of population dynamics at a yearly resolution.

If the most common *a priori* choices on the climate predictors of population fluctuations are suboptimal, then more complex models could increase predictive power. This would be the case for models that simultaneously identify the critical time window and magnitude of climatic effects that drive their system dynamics. Several such models have been proposed that automatically select the timing most relevant to population processes (van de Pol and Cockburn 2011, Ogle et al. 2015). Henceforth, we refer to these models as “antecedent effect models”. These models control for multiple comparisons, and weigh the time windows when drivers are most influential for population dynamics, dividing the climate (or any other relevant factor) of a time window (*e.g*., a year) into sub-periods of equal lengths – usually months. Such weighing can occur by estimating the importance of each monthly anomaly using weights constrained to follow a specific function (van de Pol and Cockburn 2011, Ogle et al. 2015). Alternatively, the weighing can be performed by estimating a different effect size for each monthly climate anomaly, using regularization or informative priors to minimize overfitting (Hoerl & Kennard, 1970). In cases where the timing of climatic predictors is not crucial in explaining population responses to climate, antecedent effect models would have low predictive power when compared to simpler models. Moreover, it has been suggested that, when the climatic effects on populations are small (e.g. due to their ability to buffer the effects of climate; Hilde et al. 2020), antecedent effect models would be most useful when 20 or more years of climatic data are available (Teller et al. 2016, van de Pol et al. 2016). Still, despite these potential issues, antecedent effect models provide a key advantage of being tools for exploratory analysis: the automatic estimation of the timing and magnitude of climatic effects can help generate new hypotheses.

Here, we provide an overview on antecedent effect models, we evaluate their predictive performance, and then showcase their potential for exploratory data analysis. We systematically evaluate our performance to predict the out-of-sample variation of the vital rates of survival, development, and reproduction and the overall population growth rate (λ) of 77 natural populations of 34 plant species. We compare and contrast the predictive performance of three types of antecedent effect models to that of models based on annual climatic anomalies, and null models. Here, we include null models to test the hypothesis that, in relatively small datasets, monthly or annual climatic values are inadequate predictors of variation in demographic data. We use monthly temperature and precipitation as climatic drivers, and demographic information ranging from 7-36 years from the COMPADRE Plant Matrix Database (Salguero-Gómez *et al*. 2015). Antecedent effect models were previously employed with great success on tree rings datasets (*e.g.*, Peltier et al. 2018). Here, we test these models on plant datasets for which the interest in climate effects has been steadily growing (Ehrlén et al. 2016). Our analysis goes from small, relatively common datasets to rare, long-term data that should maximize the predictive potential of antecedent effect models. We evaluate the predictive performance of these five models to address four objectives: (1) evaluate the predictive performance of each statistical model; (2) compare the ability of climate to predict among vital rates and population growth rate; (3) determine whether the predictive ability of antecedent effect models increase with temporal replication; and (4) choose a model as a case study to show the potential of antecedent effect models as exploratory data analysis tools.

## Methods

### Data and response variables

To compare the predictive performance of antecedent effect models to that of simpler models, we used long-term time series of vital rate data obtained from matrix population models (MPMs, hereafter) contained in the COMPADRE Plant Matrix Database (Salguero-Gómez et al. 2015). In this study, we ignored similar data on animals because it lacks taxonomic, temporal, and spatial replication (Salguero-Gómez et al. 2016). COMPADRE contains MPMs of natural populations where individuals have been categorized into discrete stages and their vital rates (*e.g.*, survival, development, reproduction) tracked in discrete time steps (Caswell 2001). We selected datasets from COMPADRE with at least seven contiguous years (*i.e.*, six MPMs) for a single natural population. We chose seven years as the shortest study length that allowed comparing a sufficient number of species: only half of plant demographic studies exceed four years in duration (Salguero-Gómez et al. 2015, Römer et al. 2021). This criterion resulted in 1,057 MPMs originating from 77 populations across 34 species, with an average length of 15 years (median = 10; range = 7-36), and an average number of two populations (median = 1, range=1-6). In our dataset, each study corresponded to a separate species or set of species, with the exception of *Cirsium pitcheri*. This species appeared in three separate studies, conducted on three geographically separate populations (Supplementary Online Materials Table S1). The characteristics of our species pool reflected the well-known bias of plant demographic data for temperate environments and herbaceous species (Supplementary Online Materials Table S2 (Salguero-Gómez et al. 2015, Compagnoni et al. 2021, Römer et al. 2021). Importantly, 86% our populations are eudicot, 86% herbaceous perennials, and 78% of populations are from temperate environments. We only had three shrubs, two succulent species, four species found in Mediterranean environments, and four found in deserts.

We used these MPMs to calculate the expectation of survival, development, reproduction, and the population growth rate (λ) associated with each annual MPM. By “development”, we refer to changes in life cycle stage. These four year-specific metrics refer to a population at its stable stage distribution: the distribution of discrete stages that would result if each yearly MPM’s vital rates were kept constant. We did not divide these data in vital rates of non-reproductive and reproductive plants because their year-to-year fluctuations are strongly correlated (Tredennick et al. 2018). Accordingly, our preliminary analyses on such split vital rate data showed inconspicuous differences between classes results, motivating us to lump data across size classes. We calculated survival, development, and reproduction using standard methods (Franco & Silvertown, 2004), and λ as the dominant eigenvalue of each MPM (Caswell 2001) with the R package popbio (Stubben and Milligan 2007). Here, λ > (or <) 1 corresponds to a population that is expected to grow (decline) at the stable stage distribution (Caswell, 2001). Note that we used λ instead of a population growth rate derived from year-to-year changes in population size which cannot be derived from matrix projection models. We calculated net annual vital rates by taking a weighted mean of the stage-specific vital rates, where the weights were derived from the stable stage distribution.

We used gridded and weather station climatic data to generate the climatic predictors for our statistical models. The climatic predictors in our models include mean monthly air temperature and total monthly precipitation at the location of each of our 77 populations. We obtained these data primarily from the CHELSA (Karger et al. 2017), but we also used PRISM (Daly et al. 1994) and a single Global Historical Climatology Network (GHCN) daily weather station (Menne et al. 2012) in the temporal range not covered by CHELSA. We used CHELSA data for all populations sampled between 1979 and 2013 (Supplementary Online Materials Table S1). We used PRISM data for the two populations of *Astragalus scaphoides* that extend to 2014 (Tenhumberg *et al*. 2018). We used data from weather station number USC00143527 of the GHCN for the 10 populations collected between 1938 and 1972 in Hays, Kansas, USA (Supplementary Online Materials Table S1). We used data from this weather station instead of PRISM because the quality of gridded climatic data tends to decrease with time since present. We note that CHELSA and PRISM provide gridded climatic data at a 30 arcsec (∼1 km^2^) resolution which approximate weather station climate averages well (Behnke et al. 2016). We consider this spatial resolution more than sufficient, because climate anomalies are strongly correlated spatially. For example, in North America, to observe correlations of 0.5 between the annual anomalies of two sites, one needs to travel at least 300Km, for precipitation anomalies, and 1,400 Km, for temperature anomalies (Compagnoni et al. 2024).

Our climatic predictors consisted of the monthly climatic anomalies observed in the 36 months preceding each vital rate observation (*i.e.*, preceding the end month of a given demographic projection interval). Specifically, vital rate observations describe a transition from year *t* to year *t* + 1: our models use the 36 monthly anomalies preceding year *t* + 1 as predictors. These anomalies were *z*-scores (mean = 0, SD = 1) with respect to the long-term monthly mean. For example, the anomalies for the month of December were calculated with respect to the long-term mean and standard deviation of December. We calculated long-term means and standard deviations exceeding the recommended minimum of 30 years of data (World Meteorological Organization 2017). For CHELSA and PRISM datasets, we calculated the long-term monthly mean using a 35-year period between 1979 and 2013, and 1980 and 2014, respectively. We calculated the long-term monthly mean for the 10 datasets collected before 1979 using 50 years of climate data. The difference between anomalies computed using the 50-versus 35-year mean was negligible, as their correlations were 0.996 and 0.995 for temperature and precipitation, respectively.

### Modeling overview

Using vital rates and log(λ) as response variables, we compared and contrasted the predictive performance of five models (Supplementary Online Materials: Fig. S1): the (1) Null Model (NM), with no climatic covariates, the (2) Climate Summary Model (CSM, Dalgleish et al., 2011), using yearly climate as a predictor, and three types of antecedent effect models: the (3) Weighted Means Model (WMM, van de Pol & Cockburn, 2011), the (4) Stochastic Antecedent Model (SAM, Ogle et al., 2015), and the (5) Finnish Horseshoe Model (FHM, Hastie et al., 2009). Models 1-5 are ranked by increasing flexibility, by progressively relaxing key assumptions (Supplementary Online Materials: Fig. S1). NMs are a “model of the mean” that do not account for the effect of climatic drivers. CSMs assume that the monthly anomalies observed within a specific period have all equal weight and that these anomalies affect population dynamics in the same direction. Therefore, this model uses as a predictor the mean of 12 monthly anomalies. WMM’s weigh the contribution of the 12 monthly anomalies based on a Gaussian function. The mean of this function will identify the month with the highest contribution, and all other months will have progressively lower weight (Supplementary Online Materials: Fig. S1). Like in WMMs, SAMs assume that monthly anomalies influence population dynamics in the same direction. SAMs estimate the relative weights of monthly anomalies by assigning a Dirichlet distribution prior to the weights (Supplementary Online Materials: Fig. S1; Ogle et al., 2015). Finally, FHMs fit a separate coefficient for each of the 12 monthly anomalies, shrinking these toward zero through their prior distribution. In regularized horseshoe regressions, monthly anomalies can have positive or negative effects. FHMs were not developed to model climatic effects (Piironen and Vehtari 2017), though regularization approaches have been previously employed to estimate climatic effects on vital rate information (e.g. Tredennick et al., 2016).

We chose these three antecedent effect models as they were the most useful exploratory models for our dataset. There are four types of exploratory models to estimate climate sensitivity (van de Pol and Bailey 2019): sliding windows (van de Pol et al. 2016), weighted means, regularization, and machine learning (Teller et al. 2016). We discarded sliding windows and machine learning because not easily implemented in a Bayesian framework, and because they were too complex for our small datasets. We also excluded the use of splines (Teller et al. 2016), as they are a form of regularization, and we discarded three alternatives to our WMM because of convergence issues. We provide further details on these decisions in the Supplementary Online Materials.

We fit each model type using climate data from three years: one from the year of the census, one prior to the census, and one two years before the census. We used these three years of data to account for the potential of lagged climatic effects. There is convincing evidence of lagged climatic effects in several case studies, including for three populations analyzed in this study: one population of *Cryptantha flava* (Evers et al. 2021) and two of *Astragalus scaphoides* (Tenhumberg et al., 2018). We re-fit each model using climate data from three separate years: each year corresponding to a subset of the 36 monthly anomalies preceding each demographic observation. We subdivided these 36 anomalies as belonging to the year (year *t*), one year (*t*-1), and two years (*t*-2) preceding each demographic observation. Hence, we fit five model types for each response variable. Because four of these models included climatic predictors, we refit these four models on the three years of climate data.

Below, we describe the equations of the models we tested assuming a normal response variable, log(λ). To facilitate model comparison and model convergence, we used weakly informative priors for the intercept and variance terms (Lemoine 2019). We document the priors for all of the five (Eq. 1-5) models presented below, and the prior predictive checks we ran to choose them, in the section of the Supplementary Online Materials on prior predictive checks.

Our null is a model of the mean, such that

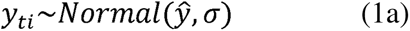

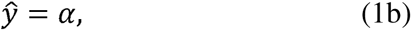

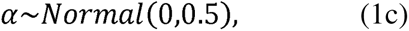

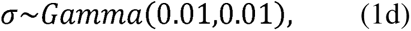

where *y_ti_*refers to a log(λ) value, *t* refers to the year of observation, *i* to a replicate within years (*i* exceeds one in datasets that are spatially replicated), (*ŷ* is the mean model prediction, σ is the standard deviation, and α is a model intercept, the mean. In the CSM, we model the response variable as a function of *x̄_t_* the mean of the 12 monthly anomalies (z-scores) over year *t,* as

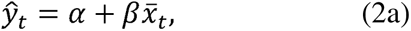

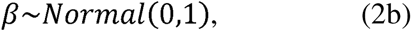

where β is the coefficient for the effect of *x̄_t_*, and *ŷ* is the mean model prediction referred to year *t*.

The first antecedent effect model is the WMM, which models the response as a function of *x_tk_*, the monthly temperature deviations for year *t* and month *k*:

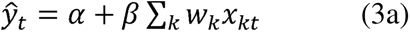

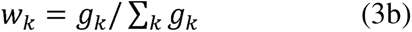

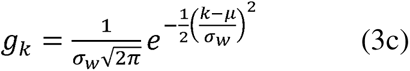

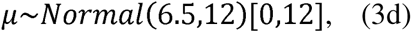

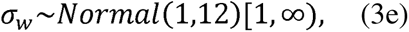

where the 12 *x_k_* monthly anomalies prior to the end of the relevant demographic year, are each weighted by *w_k_*. In turn, *w_k_* is a normalized vector of Gaussian weights, *g_k_*, with mean μ and scale σ*_w_*. These priors on the μ and σ*_w_* parameters allowed consistent convergence to these models. The prior in eq. 3d provides equal probability to μ values between 1 and 12. The prior in Eq. 3e allows values of σ*_w_* so large that the weights tend to be equal. The square brackets in Eq. 3d and 3e show the limits beyond which the potential values of these parameters are truncated. While values of μ do not extend beyond 0 and 12, σ*_w_* has no upper bound.

The SAM (Ogle et al. 2015) adds flexibility to the WMM above,

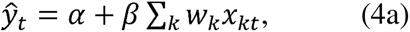

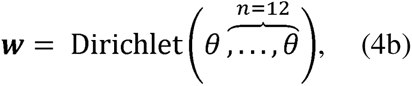

where vector ***w*** contains 12 weights that sum to 1, and *w_k_* is a single weight that refer to monthly anomaly *x_kt_* observed in month *k* and year *t*. We set the concentration parameter (*θ*) to 1, corresponding to a uniform distribution on the 12-dimensional simplex of weights. The added flexibility originates because the values in vector ***w*** do not follow a functional form (albeit they are correlated). Henceforth, we refer to the term 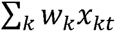, in both Eq. 3a and 4a, as the “antecedent” of year *t*.

Finally, the FHM is a multiple regression whose coefficients are shrunk via the horseshoe prior proposed by Piironen & Vehtari (2017). These models estimate a separate slope value, β*_k_*, for each *k* month. The Finnish horseshoe prior prevents overfitting by shrinking estimates towards zero. This model is fit as follows:

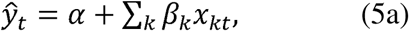

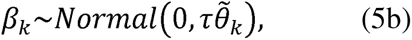

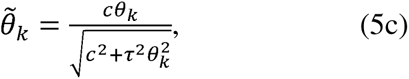

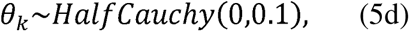

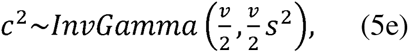

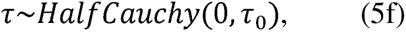

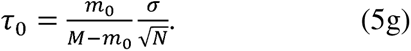

Here, 12 β*_k_* values depend on a normal distribution whose standard deviation is controlled by τ, the “global scale”, and 12 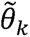 “local scales”. The global scale τ tends to shrink every monthly estimate β*_k_* toward zero. On the other hand, the 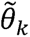 values allow the β*_k_* monthly estimates to become relatively large. In the FHM, the combination of τ and 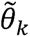 shrinks all β*_k_* coefficients toward zero, while controlling the magnitude of β*_k_* values for which there is evidence for a substantial departure from zero. We modified these models to fit survival, development, and reproduction data, which are non-normally distributed. We describe these modifications in the Supplementary Online Material.

### Model fitting

We fit models using a Hamiltonian Monte Carlo sampler via the *R* package *rstan* (Stan Development Team 2018). We fit models running four parallel chains for 16,000 iterations, with a burn-in of 4,000 iterations, and thinning by saving one out of every eight iterations. Because we had 13 alternative models, four demographic responses (log(λ) and three vital rates), two climatic predictors, and 36 datasets, we fit a total of 3,744 models. Fitting these many models made it unfeasible to check models one at a time. Therefore, we opted to perform model checks across model types, response variables, and dataset length. We checked for model convergence using four metrics. First, we flagged models with more than nine divergent transitions, models with a single parameter whose Gelman and Rubin convergence diagnostic was above 1.1, models where at least one parameter had a ratio of the Monte Carlo standard error to the standard deviation of the posterior above 0.1, and a single parameter whose effective sample size was lower than 10% of the posterior sample.

In the Supplementary Online Material, we provide the methods and results to assess the convergence of the Hamiltonian Monte Carlo samplers and assess model fit (Bayesian model checks).

### Model comparison

We compared model performance for each vital rate, climate predictor, and species combination. We performed comparisons based on the log pointwise predictive density (LPPD) (Bernardo and Smith 2009, Hooten and Hobbs 2015) calculated through a leave-one-year-out (LOYO) cross-validation (e.g. Tredennick et al., 2016). A LOYO cross-validation creates a training set by leaving out an entire year of data rather than a single data point. The model is then fit for as many years the dataset is composed of, each time leaving a different year out of the training set. The higher the value of LPPD of the held-out data, the better the predictive performance of the model. Because we refit models, each time leaving out a single year of data, we fit a total of 54,048 models. Our model selection focused on LOYO cross-validation, because we aim to predict year-level variation in vital rates. We did not perform k-fold, holdout, or exhaustive cross-validations because, despite being more conservative cross-validations, our datasets were sufficiently small as to generate convergence issues. We performed this procedure because the climate is strongly correlated among different sites in our spatially replicated datasets. In particular, 16 of our 36 datasets are spatially replicated, and their mean correlation is 0.98 (range = 0.74-1).

We compared model performance by plotting the difference in ΔLPPD — the difference in LPPD with the best model. The ΔLPPD_i_ of model *i* is calculated as LPPD*i* – max(LPPD). For each species, we plotted heat maps of the ΔLPPD_i_ values for each model, flagging the best three models. We then ordered heat maps by data length along the y-axis, and by the year of climate data (*t*, *t*-1, and *t*-2) on the x-axis.

We addressed our three objectives by testing for differences in the relative support of our models using Pseudo-Bayesian Model Averaging. We calculated model weights using Pseudo-Bayesian Model Averaging for each combination of species, response variable, and predictor (temperature or precipitation). We calculated model weights following equations 8 and 9 in Yao et al. (2018). LPPD relative weights using the same formula used for AIC weights, so that

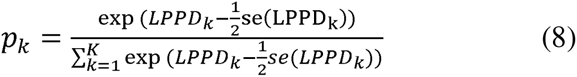

Where *p* is the LPPD relative weight, *k* refers to each model, *se* is the standard error, and *K* is 13, the total number of models tested for each combination of species, response variable, and climatic predictor. We arcsine transformed these LPPD relative weights to make them normally distributed. We then used these arcsine transformed LPPD relative weights as the response variable to address our three objectives. We addressed these objectives using graphs and average statistics. We plotted boxplots of model weights across all datasets, grouping them based on model types. This evaluated (1) the predictive performance of each of the five models. We plotted model weights of NMs to compare (2) the ability of climate to predict among vital rates and population growth rate. We also used these results to explore the role of plant functional type, ecoregion, and generation time in explaining our results. Then, we linked relative model weights against the temporal replication of each dataset to (3) verify whether such replication increased the predictive ability of antecedent effect models. We expected that replication would increase either the predictive ability of either the worst or best models. Therefore, to test this hypothesis we fit quantile regressions on these plots using the R package quantreg (Koenker 2022), from the 10^th^ to the 90^th^ quantile in increments of 10. Finally, used our model selection results to (4) select a model that showcases the potential of antecedent effect models as exploratory data analysis tools. To select the dataset of our case study, we first visually selected models with high predictive performance. Finally, we plotted these models to choose a particularly striking and biologically sensible example.

### Sensitivity analysis

We assessed the solidity of our results by running three alternative analyses: one using only the longest datasets, another fitting antecedent effect models on 36 months of climatic data, and an analysis fitting models that included both precipitation and temperature. First, we re-run analyses including only datasets that were at least 20 years long. This analysis could have increased predictive ability, because 20 years of data are considered the minimum sample size to detect a climatic signal (Teller et al. 2016, van de Pol et al. 2016a). Second, we fit our three antecedent effect models using all 36 months of climatic data. Third, we re-run all analyses including year-models using precipitation only, and models including both precipitation and temperature as predictors (using both 12 months and 36 months of data).

### Case study and exploratory analysis

We present a case study to showcase the potential of antecedent effect models as exploratory tools. We selected models with high predictive ability because they were the only ones whose model weights were unequal (for WMMs and SAMs), or whose effect sizes clearly differed from zero (for FHMs). Our case study modeled reproduction in *Astragalus tyghensis*, an endemic legume plant species with an extremely restricted range encompassing arid lands of Western North America (Thorpe and Kaye 2008). This dataset has only nine years of data but five sites, which must have substantially improved the chance to detect climatic signals (Compagnoni et al. 2024). Using this example, we first show how antecedent effect models identify monthly climatic drivers of demographic responses, and we discuss how these inferences can inform biological hypotheses. Second, we show how antecedent effect models improve model fit. We present models fit on the reproduction of *A. tyghensis* using precipitation anomalies one year prior to the last demographic observation (year t-1). This climatic predictor had high predictive power in our model comparisons. We use this case study to show for each model its fit to data and the effect sizes of monthly climatic anomalies. To show model fit to data, we plotted each raw reproduction data point against its corresponding prediction according to 200 randomly selected posterior samples. To show model fit to data, we plotted each raw reproduction data point against 200 randomly selected posterior samples predictions. To display the monthly effect sizes, we produced estimates of effect sizes suitable for a graphic comparison among CSMs, WMMs, SAMs, and FHMs. For the CSM, we divided the regression slope, β (Eq. 2a), by 12, the number of months. For the WMMs and SAMs, we multiplied the regression slope, β (Eq. 3-4a), by the 12 monthly weights, *w_k_* (Eq. 3-4b). Finally, the FHMs directly estimate 12 β*_k_* separate regression slopes. We represented these monthly effect sizes by plotting the median and the 95% credible intervals of the posterior on the y-axis, against the month number on the x-axis.

## Results

### Model comparison

Antecedent effect models were not generally useful in predicting the effects of climate, but performed well in isolated cases. When comparing LPPD weights, NMs were consistently the best performing models (Fig. 1). The variance in model weights was smaller in NMs models, while the models including climatic predictors could reach very high or very low model weights (Fig. 1). This variance in the LPPD weights suggests that estimating climate effects is advantageous for a subset of populations which are presumably particularly sensitive to climate. However, estimating climate effects can substantially decrease predictive ability in the other populations. There were no clear trends in which antecedent effect model had the best predictive performance, even if FHMs tended to perform better on log(λ) data (Fig. 1; Supplementary Online Materials: Fig. S10-13). Still, these average differences were small, potentially reflecting that these three models tend to provide similar information.

**Figure 1.**
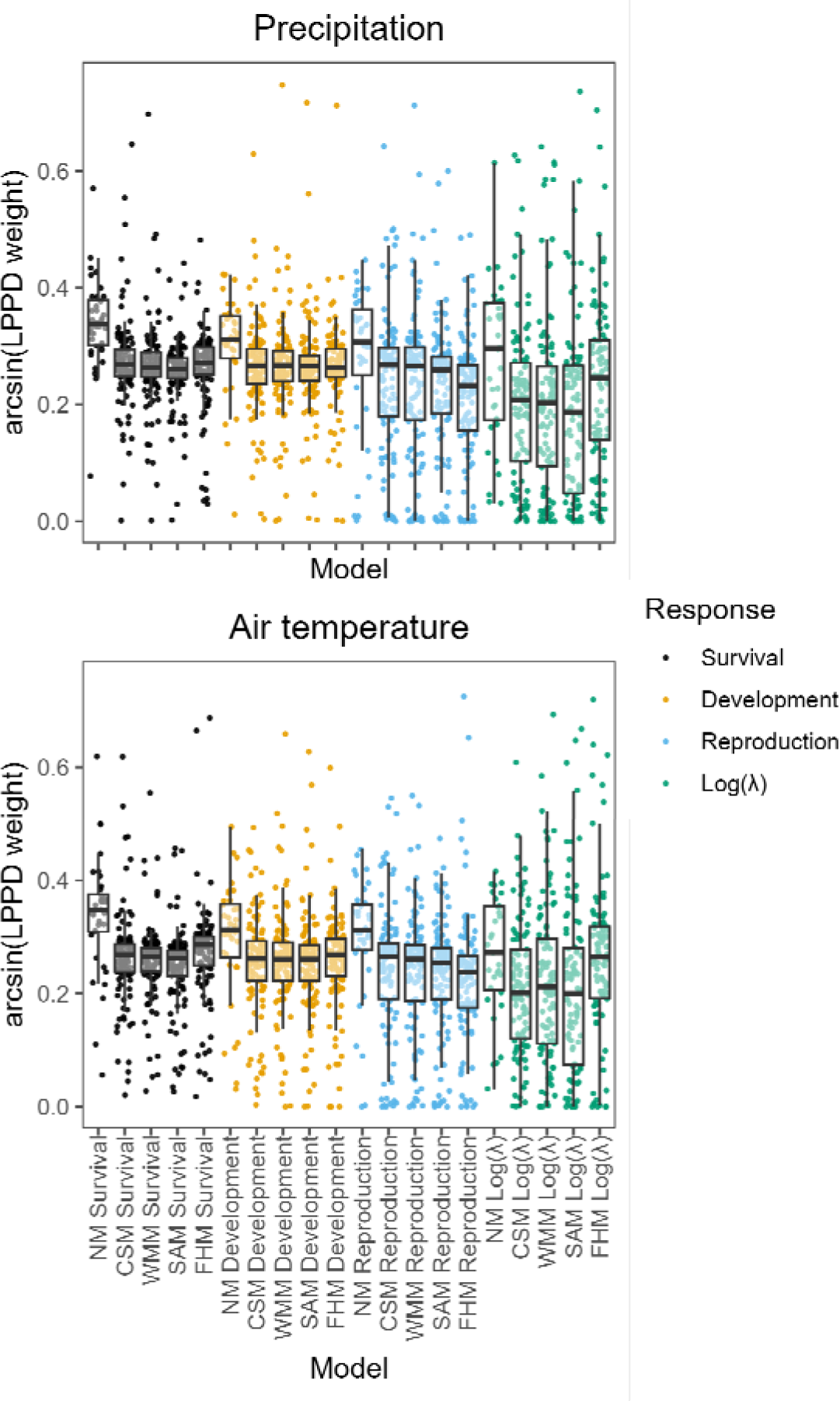
Null models (NMs) perform better than models using climate as predictor. Box and whisker plots and point clouds showing the arcsine transformed model weights for our 34 species, and across two climatic predictors, five model types, and four response variables. Panels separate models fit using precipitation or air temperature as predictor variables. The middle line of the boxplots shows the median, the upper and lower hinges delimit the first and third quartile, and the whiskers extend 1.5 times beyond the first and third quartile. Each point refers to one of the 3,744 model fits, and colors refer to the response variable of each model.

Models including climate had similar predictive performance across response variables, plant functional types, and ecoregions. On average, we found little difference in the predictive performance of response variables (Fig. 2), albeit FHMs tended to predict log(λ) better than other climate models (Fig. 1; Supplementary Online Materials: Fig. S10-11). The predictive performance was also similar across plant functional types (Supplementary Online Materials: Fig. S14), Ecoregions (Supplementary Online Materials: Fig. S15), and plant species generation time (Supplementary Online Materials: Fig. S16).

**Figure 2.**
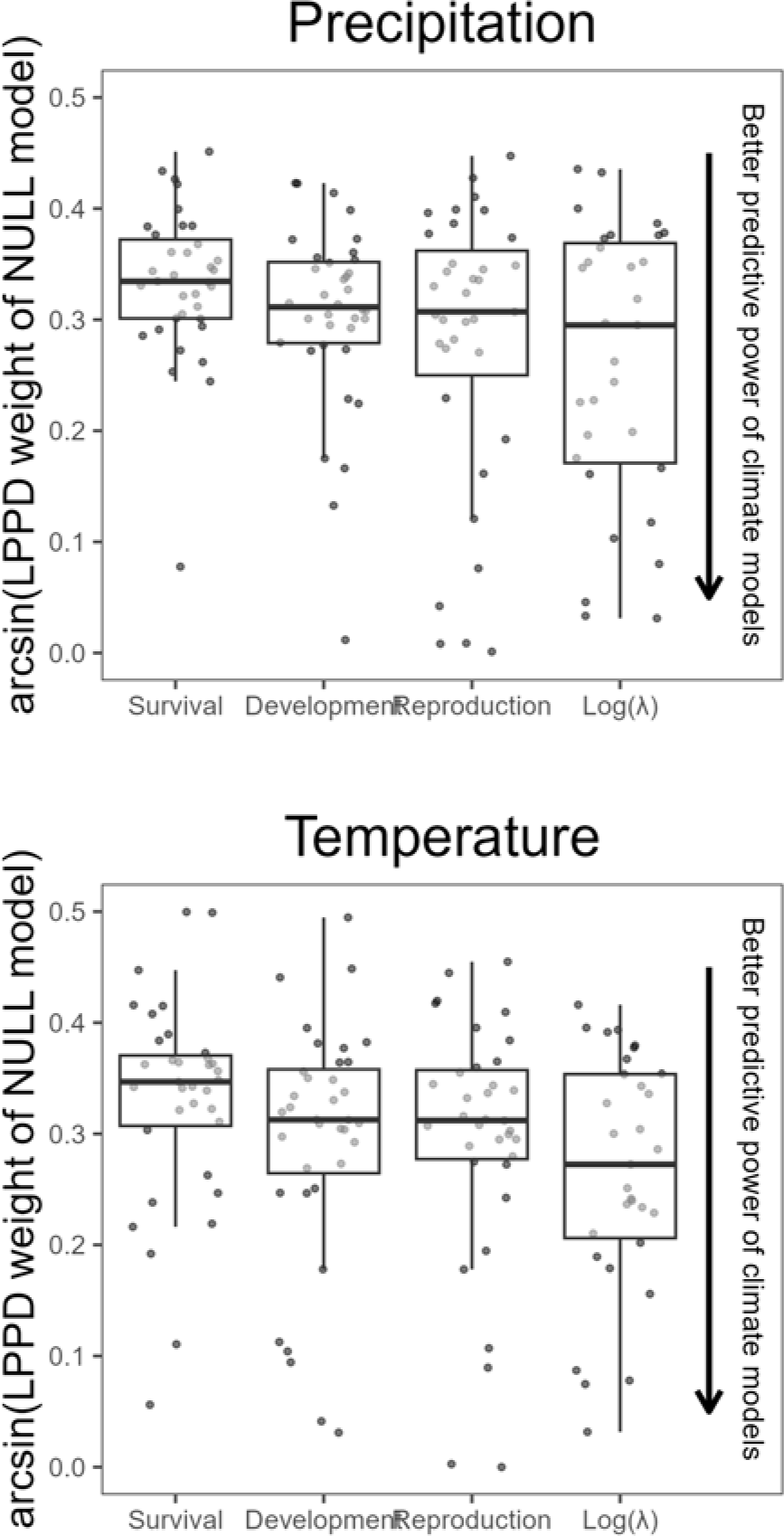
Climate has a similar predictive power across response variables. Box and whisker plots and point clouds showing the arcsine transformed model weights of the null models by climate variable, and based on the type of response variable. In this plot, lower values reflect a higher predictive ability of the models including a climatic predictor. The middle line of the boxplots shows the median, the upper and lower hinges delimit the first and third quartile, and the whiskers extend 1.5 times beyond the first and third quartile. Points represent 288 model fits: one for each species, response variable, and predictor variable (Null models were only one of the 13 models we fit on data).

The temporal replication of the studies did not improve average model performance, but it decreased the likelihood of fitting models with very low LPPD relative weights. The only quantile regression that shows a significant trend is linked to the 10^th^ quantile, for which temporal replication increases LPPD relative weights (Supplementary Online Materials: Fig. S17). This result suggests that sample size only limits the negative effects of fitting models with large numbers of parameters on predictive ability.

In our sensitivity analysis, the results of our model comparison held independently of the data and model used. First, selecting datasets longer than 20 years did not change the main results that null models have the greatest predictive performance, and that other climate models are roughly equivalent (Supplementary Online Materials: Fig. S18). Moreover, in these long-term datasets, the predictive performance of climate models was lower than that of null models across response variables (Supplementary Online Materials: Fig. S19). These results held despite these data included 36-month models. These 36-month models did not improve the predictive ability of climatic models (Supplementary Online Materials: Fig. S20-21). Finally, comparing precipitation-only models with models including precipitation plus temperature indicated that precipitation-only models tended to have better predictive ability (Supplementary Online Materials: Fig. S22-26).

### Case study and exploratory analysis

The results of antecedent effect models provided biologically interpretable information that can be used to formulate hypotheses on the demography of *A. tyghensis*. All four models show that precipitation is negatively associated with reproduction (Fig. 3). According to the WMM, the monthly anomalies preceding a demographic census tend to be the most important in driving reproduction. The SAM and FHM provide more precise insights, indicating that the most important climate windows are located two (May) and six (January) months prior to the demographic census. Thus, antecedent effect models indicate that the fecundity of this species is negatively correlated to the precipitation of the growing and dormant seasons. Therefore, in this case antecedent effect models provided information with theoretical and practical implications, even if their predictive ability was similar to the CSM (Fig. S10, Fig. 3).

**Figure 3.**
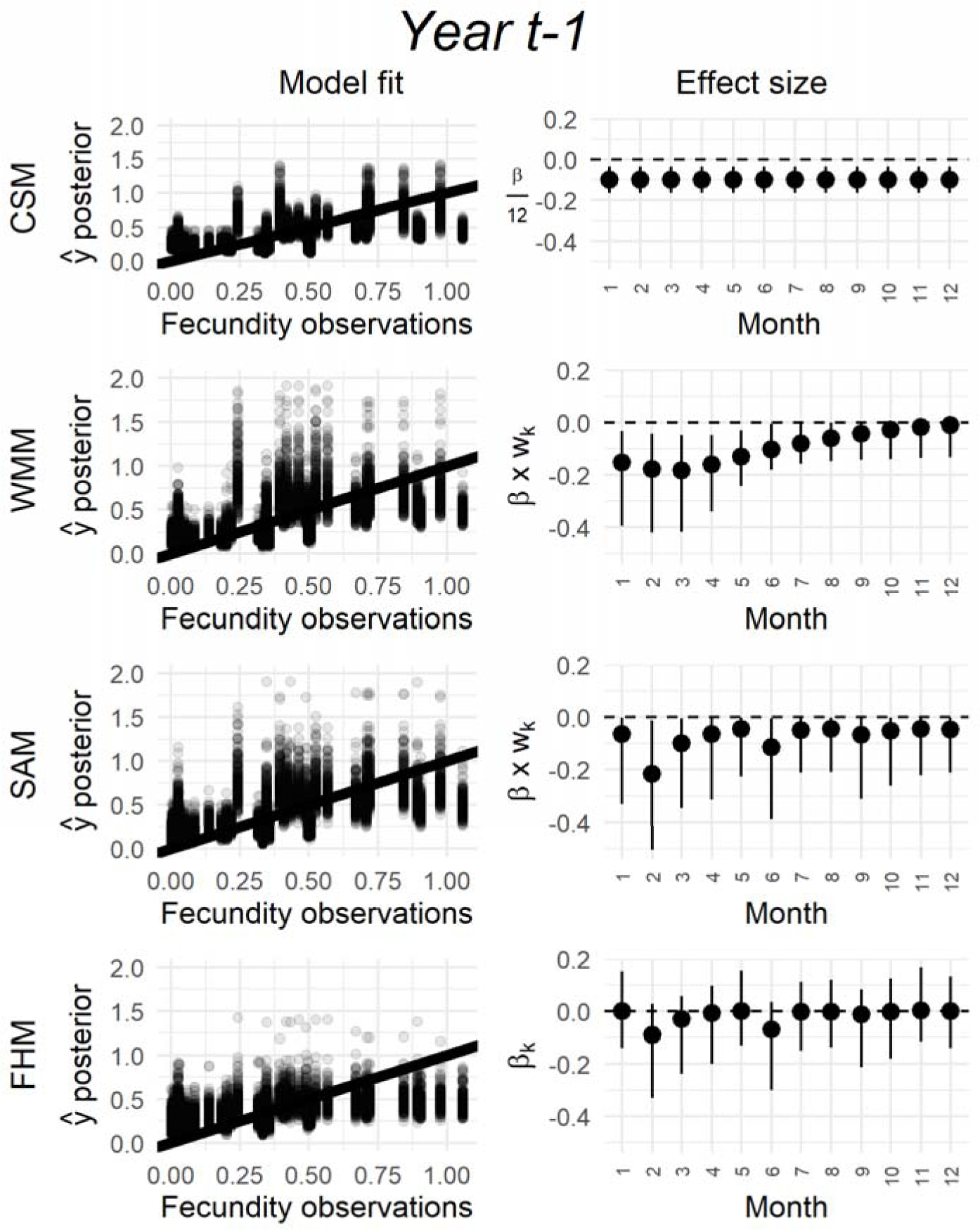
Antecedent effect models yield biologically meaningful insights. Case study showing the fits of the climate summary model (CSM), and the three antecedent effect models (WMM, SAM, FHM) to the reproduction of a natural population of the herbaceous perennial *A. tyghensis*, using precipitation anomalies two years preceding the demographic census. In each plot, columns represent, respectively, model fit and climate effect sizes. The rows refer to model type: respectively the Climate Summary Model (CSM), Weighted Mean Model (WMM), Stochastic Antecedent Model (SAM), and Finnish Horseshoe Model (FHM). In the first column, we show model fits using bivariate plots where the x-axis represents the observed reproductive values, and the y-axis represents, for each observed value, 200 randomly selected predicted values from the Bayesian posterior. In the second column, we show the effect sizes of each monthly climatic anomaly. Each monthly effect size is shown as a point range where the black circles represent the median, and the lines extend from the 2.5th to the 97.5th quantile of each posterior. In the CSM, this effect sizes correspond to the regression slope, β (Eq. 2a), divided by 12. In the WMM and SAM, the effect sizes are produced multiplying the regression slope, β (Eq. 3-4a) by the monthly weights, w_k (Eq. 3-4b). Finally, the FHM estimates a separate slope, β_k, for each monthly anomaly.

## Discussion

Predicting and understanding how climate affects the vital rates (survival, development, reproduction) of natural populations and their overall performance (*e.g.*, population growth rate, λ) is a core ecological question. By comparing the predictive performance of antecedent effect models to simpler linear models (NMs and CSMs), we found that (1) models using climate as predictor did not reliably increase our ability to predict any of the vital rates and population performance, (2) dataset length has negligible effects on predictive power, and (3) despite having similar predictive ability to CSMs, antecedent effect models provide biologically sensible insights on the timing of climatic effects. First, the hypothesize that the weak predictive ability of climate models, which is irrespective of dataset length, originates from either small climatic signals, and from low signal-to-noise ratio. A potential solution to deal with the low signal-to-noise ratio is to use individual-level data with abundant co-variate information. Second, given the generally poor predictive ability of our climate models, we conclude that on herbaceous demographic data, antecedent effect models are best used as exploratory tools for hypothesis generation. For this exploratory aim, we recommend the Finnish Horseshoe Model (FHM), because FHM has the largest flexibility.

Our initial expectation was that antecedent effect models would improve predictions, because they accommodate a variety of potential links between demography and climate (Kreyling 2010, Evers et al. 2021). This expectation was strong at least for our 12 datasets that exceeded 20 years of data, as this was the dataset length recommended for simulation studies focused on antecedent effect models (Teller et al. 2016, van de Pol et al. 2016). The predictive success of null models is a clear refutation of our initial expectation. We expected dataset length would positively affect the predictive ability of climatic variables by increasing total sample size, and by increasing the range of climatic conditions observed during the study (Compagnoni et al. 2024). For the fact that climate had small predictive power even in datasets exceeding 30 years we propose three possible explanations. First, the true effect of climate could be much smaller than what was used in previous simulation studies. For example, Teller et al. (2016) defined the climate signals they simulated as “strong”. However, subsequent analysis on the type of data mimicked by their simulations found weak climatic signals with inconsistent effects on predictive power (Tredennick et al. 2016, 2021).

Second, the signal-to-noise ratio might be quite low in aggregated data such as what we employed for this study. For example, the influential article by van de Pol et al. (2016) simulated a signal strength based on R^2^, choosing 0.2 as their lowest R^2^.Our experience with a large dataset of herbaceous plant population growth rates (Compagnoni et al. 2021) indicated a mean in-sample R^2^-squared of about 0.24. If such low signal-to-noise ratios were the norm, then accounting for confounding factors would be critical to understand and predict the effects of climate. To account for confounding factors, demographic data on individuals could facilitate accounting for co-varying factors (e.g. competition and spatial location Chu et al. 2016). Third, the way in which climate affects plants might be too complex to capture via phenomenological models applied on aggregated population data. In principle, temperature and precipitation should correlate with the main factors affecting plant processes: solar radiation, leaf temperature, soil moisture, and vapor pressure deficit (Lambers et al. 2008). Accordingly, temperature and precipitation are excellent predictors in optimal scenarios, such as tree ring studies (Fritts 2012). In these studies, much larger sample sizes belong to individuals selected to be from range positions and microsite conditions that maximize climate sensitivity (Klesse et al. 2018). On the other hand, our population dynamics data samples entire populations located at any position within a species range. Thus, our data comprises individuals and populations of varying climate sensitivity. With these type of data, unveiling climatic correlations might require sophisticated models; for example, models that approximate the microclimatic conditions at each site (e.g. Kearney and Leigh 2024).

The case study presented in this article showcases the potential of antecedent effect models for exploratory analysis and hypothesis generation. While antecedent effect models have on average similar predictive performance to CSMs, they provide granular information that is more useful scientifically. In our *A. tyghensis* case study, both the SAM and FHM showed a strong negative association between reproduction and the precipitation anomalies that occurred in the May and January of the preceding year. These results raise three hypotheses. First, such negative association with precipitation anomalies occurring a year prior to a demographic observation could result from indirect effects mediated through competition (Suttle et al. 2007, Adler et al. 2012) or physiological tradeoffs (*e.g.*, costs of reproduction, Crone et al., 2009; Miller et al., 2012). Second, the correlation with the precipitation in January, a month when the mean temperature is just above or below 0° C, could reflect the effect of snowfall. This effect is often distinct from rain (Dalgleish et al. 2011, Compagnoni and Adler 2014) and could arise in different ways: for example, because snow affects pathogens (Smull et al. 2019), it protects plants from frost (Inouye, 2000), or because snowmelt provides a large pulse of resources (Noy-Meir 1973). Third, the large effect of May precipitation could reflect a key developmental phase occurring either in May or June. All three hypotheses call for further tests to be performed via analysis of existing data or field experiments. For exploratory studies, we recommend to exploit the flexibility of FHMs. FHMs usually provide qualitatively similar results to the SAMs, but with the added benefit of simultaneously identifying positive and negative climatic signals. While it is unlikely for climate signals to go in opposite directions, we did observe such cases in some of our model fits. It is perhaps because of this flexibility that FHMs outperform WMMs and SAMs when predicting log(λ) (Figure 1).

Our experience building and comparing antecedent effect models informs our suggestions on dataset length, hyperparameter priors, and emphasizes the role of prior predictive checks. While most of our models had no problems converging, problems occasionally arose when datasets had less than about 20 years of data (Fig. S5-6). Therefore, in our case the often recommended dataset length of 20 years (Teller et al. 2016, van de Pol and Bailey 2019) improved model convergence rather than model predictive ability. The choice of hyperparameter priors has strong effects on model fitting and performance. In weighted mean models (WMMs), truncated hyperparameter priors (Eq. 3d-e) prevented the sampler from exploring unrealistic parameter values, greatly facilitating model convergence. In SAMs, we chose a vector of ones as the Dirichlet concentration parameter (Eq. 4b) to not favor any particular lag, and to ensure that prior weights had a uniform distribution. A much lower number than 1 would favor models with a single non-zero weight. In FHMs, we suggest using a standard deviation of 0.1 for the prior on monthly effect sizes (Eq. 5d). Using a standard deviation of 1 (*e.g.*, Piironen & Vehtari, 2017) generated unrealistically high variation in our prior predictive checks (Supplementary Online Materials: prior predictive checks). More generally, prior predictive checks were essential because our initial prior choices produced very different variance in predictions among model types and response variables. Moreover, priors generating unrealistically high variance had model convergence issues and lower predictive performance.

These results justify the continued data and conceptual efforts to understand and predict the effects of climatic drivers on the vital rates of herbaceous plants. These effects of climate are complex, and antecedent effect models could contribute to understand this complexity as a hypothesis generation tool. However, improvements in predictive ability will rely on datasets with larger replication, especially temporal replication. Moreover, conceptual breakthroughs such as those regarding the importance of indirect effects (Clark et al., 2021; Hacket Pain et al., 2018), might also prove useful.

## Data availability statement

The data and reproducible code for the analyses connected to this study are available at https://zenodo.org/records/7839199.

## Acknowledgements

AC and TMK were supported by the Alexander von Humboldt Foundation under the framework of the Humboldt Professorship, and by the Helmholtz Recruitment Initiative of the Helmholtz Association (both to TMK); RSG was supported by NERC (NE/M018458/1 and NE/X103766/1). This work emanated from research discussions during the sAPROPOS (Analysis of PROjections of POpulationS) working group led by RSG and TMK at the German Centre for Integrative Biodiveristy Research (iDiv), which is funded by the German Research Foundation (FZT-118-202548816). We thank the Max Planck Institute for Demographic Research for the support of the COMPADRE Plant Matrix Database and the hundreds of plant population ecologists who have contributed with demographic data to COMPADRE. We thank B. Teller and P. Barks for discussions in the early phases of this project.

## Author Contributions

**Aldo Compagnoni**: Conceptualization (Equal); Data curation (Equal); Formal analysis (Lead); Methodology (Lead); Software (Lead); Validation (Lead); Visualization (Lead); Writing – original draft (Lead); Writing – review & editing (Equal). **Dylan Z. Childs**: Conceptualization (Supporting); Data curation (Equal); Formal analysis (Supporting); Methodology (Supporting); Project administration (Supporting); Software (Supporting); Supervision (Supporting); Validation (Supporting); Writing – review & editing (Equal). **Tiffany M Knight**: Data curation (Equal); Funding acquisition (Lead); Resources (Supporting); Supervision (Supporting); Writing – review & editing (Equal). **Roberto Salguero-Gómez**: Conceptualization (Equal); Data curation (Equal); Formal analysis (Supporting); Funding acquisition (Supporting); Methodology (Supporting); Project administration (Lead); Supervision (Lead); Writing – review & editing (Equal).

