## Supplementary online material for "Antecedent effect models as an exploratory tool to link climate drivers to herbaceous perennial population dynamics data"

Supplementary Table S1. Plant Populations examined in this study. We include the name of species in the COMPADRE Plant Matrix Database, the accepted species name, and the species labels identify species consistently with figures S12 and S13. For each species, we also present the name of the populations at which it was censused, the starting and ending year of census, and the type of climate data we used to fit statistical models. Climate data was obtained from either CHELSA (Karger et al., 2017), PRISM (Daly et al., 1994), or Global Historical Climatology Network (GHCN; Menne et al., 2012).

| **Species name in COMPADRE** | **Population name** | **Start year** | **End year** | **Climate data** | **Accepted species name** | **Species labels** |
|---|---|---|---|---|---|---|
| Actaea_spicata | Site A | 1992 | 1998 | CHELSA | *Actaea spicata* | Actaea spicata |
| Actaea_spicata | Site B | 1992 | 1998 | CHELSA | *Actaea spicata* | Actaea spicata |
| Astragalus_scaphoides_6 | McDevitt Creek | 1988 | 2014 | PRISM | *Astragalus scaphoides* | Astragalus scaphoides |
| Astragalus_scaphoides_6 | Sheep Corral Gulch | 1988 | 2014 | PRISM | *Astragalus scaphoides* | Astragalus scaphoides |
| Astragalus_tyghensis | Site 10 | 1991 | 2000 | CHELSA | *Astragalus tyghensis* | Astragalus tyghensis |
| Astragalus_tyghensis | Site 13 | 1991 | 2000 | CHELSA | *Astragalus tyghensis* | Astragalus tyghensis |
| Astragalus_tyghensis | Site 25 | 1991 | 2000 | CHELSA | *Astragalus tyghensis* | Astragalus tyghensis |
| Astragalus_tyghensis | Site 4 | 1991 | 2000 | CHELSA | *Astragalus tyghensis* | Astragalus tyghensis |
| Astragalus_tyghensis | Site 41 | 1991 | 2000 | CHELSA | *Astragalus tyghensis* | Astragalus tyghensis |
| Arabis_fecunda | Charleys Gulch | 1987 | 1993 | CHELSA | *Boechera fecunda* | Arabis fecunda |
| Brassica_insularis | Calcina | 2000 | 2009 | CHELSA | *Brassica insularis* | Brassica insularis |
| Brassica_insularis | Corbaghiola | 2000 | 2009 | CHELSA | *Brassica insularis* | Brassica insularis |
| Brassica_insularis | Inzecca | 2000 | 2009 | CHELSA | *Brassica insularis* | Brassica insularis |
| Brassica_insularis | Teghime | 2000 | 2009 | CHELSA | *Brassica insularis* | Brassica insularis |
| Cirsium_pitcheri_4 | CiPi 1 | 1988 | 1998 | CHELSA | *Cirsium pitcheri* | Cirsium pitcheri (2) |
| Cirsium_pitcheri_4 | CiPi 2 | 1988 | 1998 | CHELSA | *Cirsium pitcheri* | Cirsium pitcheri (2) |
| Cirsium_pitcheri_4 | CiPi 3 | 1988 | 1998 | CHELSA | *Cirsium pitcheri* | Cirsium pitcheri (2) |
| Cirsium_pitcheri_6 | Illinois Beach State Park | 1997 | 2004 | CHELSA | *Cirsium pitcheri* | Cirsium pitcheri (3) |
| Cirsium_pitcheri_8 | Wilderness State Park | 1995 | 2013 | CHELSA | *Cirsium pitcheri* | Cirsium pitcheri (1) |
| Cirsium_undulatum | Hays | 1942 | 1972 | GHCN | *Cirsium undulatum* | Cirsium undulatum |
| Cypripedium_calceolus | Oparzelisko | 1991 | 1999 | CHELSA | *Cypripedium calceolus* | Cypripedium calceolus |

| | | | | | | |
|---|---|---|---|---|---|---|
| Cypripedium_calceolus | Pogorzaly | 1991 | 1999 | CHELSA | *Cypripedium calceolus* | Cypripedium calceolus |
| Cypripedium_fasciculatum | Region 3 | 1999 | 2007 | CHELSA | *Cypripedium fasciculatum* | Cypripedium fasciculatum |
| Cypripedium_fasciculatum | Region 4 | 1999 | 2007 | CHELSA | *Cypripedium fasciculatum* | Cypripedium fasciculatum |
| Cypripedium_fasciculatum | Region 5 | 1999 | 2007 | CHELSA | *Cypripedium fasciculatum* | Cypripedium fasciculatum |
| Dicerandra_frutescens | Fireline Edge 1 (Pop 0) | 1988 | 1995 | CHELSA | *Dicerandra frutescens* | Dicerandra frutescens |
| Dicerandra_frutescens | Oak-hickory scrub 1 (Pop 10) | 1988 | 1995 | CHELSA | *Dicerandra frutescens* | Dicerandra frutescens |
| Dicerandra_frutescens | Oak-hickory scrub 3 (Pop 19) | 1990 | 2000 | CHELSA | *Dicerandra frutescens* | Dicerandra frutescens |
| Dicerandra_frutescens | Sand pine scrub 1 (Pop 2) | 1988 | 2000 | CHELSA | *Dicerandra frutescens* | Dicerandra frutescens |
| Dicerandra_frutescens | Sand pine scrub 2 (Pop 4) | 1988 | 1997 | CHELSA | *Dicerandra frutescens* | Dicerandra frutescens |
| Dicerandra_frutescens | Yard Edge (Pop 24) | 1991 | 2000 | CHELSA | *Dicerandra frutescens* | Dicerandra frutescens |
| Echinacea_angustifolia | Hays | 1938 | 1971 | GHCN | *Echinacea angustifolia* | Echinacea angustifolia |
| Eriogonum_longifolium_var._gnaphalifolium_2 | Unburned | 1990 | 1998 | CHELSA | *Eriogonum longifolium gnaphalifolium* | Eriogonum_longifolium... |
| Eryngium_alpinum | BER | 2001 | 2010 | CHELSA | *Eryngium alpinum* | Eryngium alpinum |
| Eryngium_alpinum | DES | 2001 | 2010 | CHELSA | *Eryngium alpinum* | Eryngium alpinum |
| Eryngium_alpinum | PRA | 2001 | 2010 | CHELSA | *Eryngium alpinum* | Eryngium alpinum |
| Eryngium_alpinum | PRB | 2001 | 2010 | CHELSA | *Eryngium alpinum* | Eryngium alpinum |
| Eryngium_alpinum | PRC | 2001 | 2010 | CHELSA | *Eryngium alpinum* | Eryngium alpinum |
| Eryngium_alpinum | PRD | 2001 | 2010 | CHELSA | *Eryngium alpinum* | Eryngium alpinum |
| Eryngium_cuneifolium | 72 | 1990 | 1998 | CHELSA | *Eryngium cuneifolium* | Eryngium cuneifolium |
| Eryngium_cuneifolium | 85 | 1990 | 1999 | CHELSA | *Eryngium cuneifolium* | Eryngium cuneifolium |
| Eryngium_cuneifolium | 91 | 1990 | 1998 | CHELSA | *Eryngium cuneifolium* | Eryngium cuneifolium |

| | | | | | | |
|---|---|---|---|---|---|---|
| Helianthemum_juliae | Teide National Park | 1992 | 2001 | CHELSA | *Helianthemum juliae* | Helianthemum juliae |
| Horkelia_congesta | Long Tom ACEC | 1993 | 1999 | CHELSA | *Horkelia congesta* | Horkelia congesta |
| Kosteletzkya_pentacarpos | Ricarda lagoon | 1996 | 2004 | CHELSA | *Kosteletzkya pentacarpos* | Kosteletzkya pentacarpos |
| Lomatium_bradshawii | Long Tom | 1988 | 1997 | CHELSA | *Lomatium bradshawii* | Lomatium bradshawii |
| Opuntia_macrorhiza_2 | Plot 52 | 1999 | 2005 | CHELSA | *Opuntia macrorhiza* | Opuntia macrorhiza |
| Opuntia_macrorhiza_2 | Plot 57 | 1999 | 2005 | CHELSA | *Opuntia macrorhiza* | Opuntia macrorhiza |
| Orchis_purpurea | Light Environment 1 | 2002 | 2008 | CHELSA | *Orchis purpurea* | Orchis purpurea |
| Orchis_purpurea | Light Environment 2 | 2002 | 2008 | CHELSA | *Orchis purpurea* | Orchis purpurea |
| Orchis_purpurea | Light Environment 3 | 2002 | 2008 | CHELSA | *Orchis purpurea* | Orchis purpurea |
| Orchis_purpurea | Shaded site 1 | 2002 | 2008 | CHELSA | *Orchis purpurea* | Orchis purpurea |
| Orchis_purpurea | Shaded site 2 | 2002 | 2008 | CHELSA | *Orchis purpurea* | Orchis purpurea |
| Orchis_purpurea | Shaded site 3 | 2002 | 2008 | CHELSA | *Orchis purpurea* | Orchis purpurea |
| Paronychia_jamesii | Hays | 1937 | 1972 | GHCN | *Paronychia jamesii* | Paronychia jamesii |
| Pediocactus_bradyi | Badger Creek | 1988 | 2010 | CHELSA | *Pediocactus bradyi* | Pediocactus bradyi |
| Pediocactus_bradyi | North Canyon East | 1988 | 2010 | CHELSA | *Pediocactus bradyi* | Pediocactus bradyi |
| Pediocactus_bradyi | North Canyon West | 1988 | 2010 | CHELSA | *Pediocactus bradyi* | Pediocactus bradyi |
| Pediocactus_bradyi | Soap Creek | 1988 | 2010 | CHELSA | *Pediocactus bradyi* | Pediocactus bradyi |
| Lesquerella_ovalifolia | Hays | 1937 | 1972 | GHCN | *Physaria ovalifolia* | Lesquerella ovalifolia |
| Psoralea_tenuiflora | Hays | 1938 | 1972 | GHCN | *Psoralea tenuiflora* | Psoralea tenuiflora |
| Purshia_subintegra | Dry | 1996 | 2003 | CHELSA | *Purshia subintegra* | Purshia subintegra |
| Purshia_subintegra | Moist | 1996 | 2003 | CHELSA | *Purshia subintegra* | Purshia subintegra |
| Haplopappus_radiatus | 1in | 1991 | 2000 | CHELSA | *Pyrrocoma radiata* | Haplopappus radiatus |
| Haplopappus_radiatus | 2in | 1991 | 2000 | CHELSA | *Pyrrocoma radiata* | Haplopappus radiatus |
| Haplopappus_radiatus | 3in | 1991 | 2000 | CHELSA | *Pyrrocoma radiata* | Haplopappus radiatus |
| Haplopappus_radiatus | 4in | 1991 | 2000 | CHELSA | Pyrrocoma radiata | Haplopappus radiatus |

| | | | | | | |
|---|---|---|---|---|---|---|
| Haplopappus_radiatus | 5in | 1991 | 2000 | CHELSA | Pyrrocoma radiata | Haplopappus radiatus |
| Ratibida_columnifera | Hays | 1941 | 1972 | GHCN | Ratibida columnifera | Ratibida columnifera |
| Silene_spaldingii | Eureka | 1987 | 1999 | CHELSA | Silene spaldingii | Silene spaldingii |
| Solidago_mollis | Hays | 1941 | 1972 | GHCN | Solidago mollis | Solidago mollis |
| Sphaeralcea_coccinea | Hays | 1937 | 1972 | GHCN | Sphaeralcea coccinea | Sphaeralcea coccinea |
| Hedyotis_nigricans | Hays | 1943 | 1972 | GHCN | Stenaria nigricans | Hedyotis nigricans |
| Thelesperma_megapotamicum | Hays | 1938 | 1971 | GHCN | Thelesperma megapotamicum | Thelesperma megapotamicum |
| Trillium_ovatum | Big Creek, Bitterroot Valley (Bitterroot National Forest) | 2003 | 2009 | CHELSA | Trillium ovatum | Trillium ovatum |
| Trillium_ovatum | Grant Creek, Missoula Valley (Lolo National Forest) | 2003 | 2009 | CHELSA | Trillium ovatum | Trillium ovatum |
| Cryptantha_flava_2 | Redfleet State Park | 1997 | 2012 | CHELSA | Cryptantha flava | Cryptantha flava |

Daly, C., Neilson, R. P., & Phillips, D. L. (1994). A Statistical-Topographic Model for Mapping Climatological Precipitation over Mountainous Terrain. *Journal of Applied Meteorology*, *33*(2), 140–158. https://doi.org/10.1175/1520-0450(1994)033<0140:ASTMFM>2.0.CO;2

Karger, D. N., Conrad, O., Böhner, J., Kawohl, T., Kreft, H., Soria-Auza, R. W., Zimmermann, N. E., Linder, H. P., & Kessler, M. (2017). Climatologies at high resolution for the earth's land surface areas. *Scientific Data*, *4*(1), 1–20. https://doi.org/10.1038/sdata.2017.122

Menne, M. J., Durre, I., Vose, R. S., Gleason, B. E., & Houston, T. G. (2012). An Overview of the Global Historical Climatology Network-Daily Database. *Journal of Atmospheric and Oceanic Technology*, *29*(7), 897–910. https://doi.org/10.1175/JTECH-D-11-00103.1

Supplementary Table S2. Plant functional types of study species, and ecoregions and mean climate of the study populations. Species labels identify species consistently with figures S12 and S13. Ecoregion refer to the classification by Olson et al. (2001). The acronyms of the last three column names refer to mean annual temperature (MAT), mean annual precipitation (MAP), and water availability index (WAI), which is given by MAP minus potential evapotranspiration.

| Species label | Population | Organism Type | Dicot/ Monoc | Country | Continent | Ecoregion | MAT | MAP | WAI |
|---|---|---|---|---|---|---|---|---|---|
| Actaea spicata | Site A | Herbaceous perennial | Eudicot | SWE | Europe | TBM | 6.9 | 584.7 | -17.4 |
| Actaea spicata | Site B | Herbaceous perennial | Eudicot | SWE | Europe | TBM | 6.6 | 612.1 | -6.6 |
| Arabis fecunda | Charleys Gulch | Herbaceous perennial | Eudicot | USA | N America | TGS | 5.7 | 454 | -36.6 |
| Astragalus scaphoides | McDevitt Creek | Herbaceous perennial | Eudicot | USA | N America | TCF | 3.6 | 324.9 | -54.4 |
| Astragalus scaphoides | Sheep Corral Gulch | Herbaceous perennial | Eudicot | USA | N America | TCF | 4.1 | 220.4 | -66.5 |
| Astragalus tyghensis | Site 10 | Herbaceous perennial | Eudicot | USA | N America | TGS | 10.1 | 577 | -43 |
| Astragalus tyghensis | Site 13 | Herbaceous perennial | Eudicot | USA | N America | TGS | 10.1 | 587.7 | -41.8 |
| Astragalus tyghensis | Site 25 | Herbaceous perennial | Eudicot | USA | N America | TGS | 11.1 | 381.3 | -53.1 |
| Astragalus tyghensis | Site 4 | Herbaceous perennial | Eudicot | USA | N America | TGS | 10.1 | 577 | -43 |
| Astragalus tyghensis | Site 41 | Herbaceous perennial | Eudicot | USA | N America | TGS | 11.7 | 341.7 | -54 |
| Brassica insularis | Calcina | Herbaceous perennial | Eudicot | FRA | Europe | MED | 16.1 | 743.5 | -46.5 |
| Brassica insularis | Corbaghiola | Herbaceous perennial | Eudicot | FRA | Europe | MED | 11.6 | 1000.9 | -12.5 |

| | | | | | | | | | |
|---|---|---|---|---|---|---|---|---|---|
| Brassica insularis | Inzecca | Herbaceous perennial | Eudicot | FRA | Europe | MED | 13.6 | 926.1 | -10 |
| Brassica insularis | Teghime | Herbaceous perennial | Eudicot | FRA | Europe | MED | 11.9 | 820.8 | -26.7 |
| Cirsium pitcheri (1) | Wilderness State Park | Herbaceous perennial | Eudicot | USA | N America | TBM | 6.6 | 867 | -2.5 |
| Cirsium pitcheri (2) | CiPi 1 | Herbaceous perennial | Eudicot | USA | N America | TGS | 10.4 | 1145.6 | 3.8 |
| Cirsium pitcheri (2) | CiPi 2 | Herbaceous perennial | Eudicot | USA | N America | TGS | 10.4 | 1161 | 9.9 |
| Cirsium pitcheri (2) | CiPi 3 | Herbaceous perennial | Eudicot | USA | N America | TGS | 10.4 | 1112.6 | 1.6 |
| Cirsium pitcheri (3) | Illinois Beach State Park | Herbaceous perennial | Eudicot | USA | N America | TGS | 9.5 | 962.5 | -17.8 |
| Cirsium undulatum | Hays | Herbaceous perennial | Eudicot | USA | N America | TGS | 13 | 693.4 | -58 |
| Cryptantha flava | Redfleet State Park | Herbaceous perennial | Eudicot | USA | N America | DES | 9.5 | 250.6 | -72.1 |
| Cypripedium calceolus | Oparzelisko | Herbaceous perennial | Monocot | POL | Europe | TBM | 7.7 | 568.7 | -14.4 |
| Cypripedium calceolus | Pogorzaly | Herbaceous perennial | Monocot | POL | Europe | TBM | 7.7 | 568.7 | -14.4 |
| Cypripedium fasciculatum | Region 3 | Herbaceous perennial | Monocot | USA | N America | TCF | 11.8 | 1470 | 33.9 |
| Cypripedium fasciculatum | Region 4 | Herbaceous perennial | Monocot | USA | N America | TCF | 12.8 | 815.9 | -20.9 |
| Cypripedium fasciculatum | Region 5 | Herbaceous perennial | Monocot | USA | N America | TCF | 10.3 | 794.3 | -20.5 |
| Dicerandra frutescens | Fireline Edge 1 (Pop 0) | Herbaceous perennial | Eudicot | USA | N America | TCF | 22.9 | 1414.2 | -20.8 |

| | | | | | | | | | |
|---|---|---|---|---|---|---|---|---|---|
| Dicerandra frutescens | Oak-hickory scrub 1 (Pop 10) | Herbaceous perennial | Eudicot | USA | N America | TCF | 22.9 | 1414.5 | -20.9 |
| Dicerandra frutescens | Oak-hickory scrub 3 (Pop 19) | Herbaceous perennial | Eudicot | USA | N America | TCF | 22.9 | 1414.4 | -20.8 |
| Dicerandra frutescens | Sand pine scrub 1 (Pop 2) | Herbaceous perennial | Eudicot | USA | N America | TCF | 22.9 | 1414.4 | -20.8 |
| Dicerandra frutescens | Sand pine scrub 2 (Pop 4) | Herbaceous perennial | Eudicot | USA | N America | TCF | 22.9 | 1414.4 | -20.8 |
| Dicerandra frutescens | Yard Edge (Pop 24) | Herbaceous perennial | Eudicot | USA | N America | TCF | 22.9 | 1414.4 | -20.8 |
| Echinacea angustifolia | Hays | Herbaceous perennial | Eudicot | USA | N America | TGS | 13 | 693.4 | -58 |
| Eriogonum_longifolium... | Unburned | Herbaceous perennial | Eudicot | USA | N America | TCF | 22.9 | 1419.3 | -21.5 |
| Eryngium alpinum | BER | Herbaceous perennial | Eudicot | FRA | Europe | TBM | 4.9 | 1262.5 | 40.6 |
| Eryngium alpinum | DES | Herbaceous perennial | Eudicot | FRA | Europe | TBM | 4.9 | 1262.5 | 40.6 |
| Eryngium alpinum | PRA | Herbaceous perennial | Eudicot | FRA | Europe | TBM | 2.7 | 1212.6 | 40.3 |
| Eryngium alpinum | PRB | Herbaceous perennial | Eudicot | FRA | Europe | TBM | 2.7 | 1212.6 | 40.3 |
| Eryngium alpinum | PRC | Herbaceous perennial | Eudicot | FRA | Europe | TBM | 4.9 | 1262.5 | 40.6 |
| Eryngium alpinum | PRD | Herbaceous perennial | Eudicot | FRA | Europe | TBM | 4.9 | 1262.5 | 40.6 |

| | | | | | | | | | |
|---|---|---|---|---|---|---|---|---|---|
| Eryngium cuneifolium | 72 | Herbaceous perennial | Eudicot | USA | N America | TCF | 22.9 | 1409.1 | -23.6 |
| Eryngium cuneifolium | 85 | Herbaceous perennial | Eudicot | USA | N America | TCF | 22.9 | 1409.1 | -23.6 |
| Eryngium cuneifolium | 91 | Herbaceous perennial | Eudicot | USA | N America | TCF | 22.9 | 1409.1 | -23.6 |
| Haplopappus radiatus | 1in | Herbaceous perennial | Eudicot | USA | N America | DES | 10.9 | 306.6 | -74.5 |
| Haplopappus radiatus | 2in | Herbaceous perennial | Eudicot | USA | N America | DES | 10.9 | 306.6 | -74.5 |
| Haplopappus radiatus | 3in | Herbaceous perennial | Eudicot | USA | N America | DES | 10.9 | 306.6 | -74.5 |
| Haplopappus radiatus | 4in | Herbaceous perennial | Eudicot | USA | N America | DES | 10.9 | 306.6 | -74.5 |
| Haplopappus radiatus | 5in | Herbaceous perennial | Eudicot | USA | N America | DES | 10.9 | 306.6 | -74.5 |
| Hedyotis nigricans | Hays | Herbaceous perennial | Eudicot | USA | N America | TGS | 13 | 693.4 | -58 |
| Helianthemum juliae | Teide National Park | Shrub | Eudicot | ESP | Europe | MED | 10.4 | 467 | -95.1 |
| Horkelia congesta | Long Tom ACEC | Herbaceous perennial | Eudicot | USA | N America | TGS | 11.7 | 1472.4 | 36.7 |
| Kosteletzkya pentacarpos | Ricarda lagoon | Herbaceous perennial | Eudicot | ESP | Europe | MED | 16.5 | 588.7 | -53 |
| Lesquerella ovalifolia | Hays | Herbaceous perennial | Eudicot | USA | N America | TGS | 13 | 693.4 | -58 |
| Lomatium bradshawii | Long Tom | Herbaceous perennial | Eudicot | USA | N America | TCF | 11.6 | 1526.2 | 40.5 |
| Opuntia macrorhiza | Plot 52 | Succulent | Eudicot | USA | N America | TCF | 8.3 | 480.2 | -55.3 |
| Opuntia macrorhiza | Plot 57 | Succulent | Eudicot | USA | N America | TCF | 8.4 | 466.7 | -51.4 |

| | | | | | | | | | |
|---|---|---|---|---|---|---|---|---|---|
| Orchis purpurea | Light Environment 1 | Herbaceous perennial | Monocot | BEL | Europe | TBM | 10.2 | 893 | 1.9 |
| Orchis purpurea | Light Environment 2 | Herbaceous perennial | Monocot | BEL | Europe | TBM | 10.2 | 893 | 1.9 |
| Orchis purpurea | Light Environment 3 | Herbaceous perennial | Monocot | BEL | Europe | TBM | 10.2 | 893 | 1.9 |
| Orchis purpurea | Shaded site 1 | Herbaceous perennial | Monocot | BEL | Europe | TBM | 10.2 | 893 | 1.9 |
| Orchis purpurea | Shaded site 2 | Herbaceous perennial | Monocot | BEL | Europe | TBM | 10.2 | 893 | 1.9 |
| Orchis purpurea | Shaded site 3 | Herbaceous perennial | Monocot | BEL | Europe | TBM | 10.2 | 893 | 1.9 |
| Paronychia jamesii | Hays | Herbaceous perennial | Eudicot | USA | N America | TGS | 13 | 693.4 | -58 |
| Pediocactus bradyi | Badger Creek | Succulent | Eudicot | USA | N America | DES | 10.2 | 763.7 | -27.7 |
| Pediocactus bradyi | North Canyon East | Succulent | Eudicot | USA | N America | DES | 8.5 | 529.3 | -77.6 |
| Pediocactus bradyi | North Canyon West | Succulent | Eudicot | USA | N America | DES | 8.5 | 529.3 | -77.6 |
| Pediocactus bradyi | Soap Creek | Succulent | Eudicot | USA | N America | DES | 16.4 | 217.4 | -101.4 |
| Psoralea tenuiflora | Hays | Herbaceous perennial | Eudicot | USA | N America | TGS | 12.6 | 621.6 | -59.4 |
| Purshia subintegra | Dry | Shrub | Eudicot | USA | N America | DES | 17 | 364.7 | -86.6 |
| Purshia subintegra | Moist | Shrub | Eudicot | USA | N America | DES | 17 | 364.7 | -86.6 |
| Ratibida columnifera | Hays | Herbaceous perennial | Eudicot | USA | N America | TGS | 12.6 | 621.6 | -59.4 |

Olson, D. M., Dinerstein, E., Wikramanayake, E. D., Burgess, N. D., Powell, G. V. N., Underwood, E. C., D'amico, J. A., Itoua, I., Strand, H. E., Morrison, J. C., Loucks, C. J., Allnutt, T. F., Ricketts, T. H., Kura, Y., Lamoreux, J. F., Wettengel, W. W., Hedao, P., & Kassem, K. R. (2001). Terrestrial Ecoregions of the World: A New Map of Life on Earth: A new global map of terrestrial ecoregions provides an innovative tool for conserving biodiversity. *BioScience*, *51*(11), 933–938. https://doi.org/10.1641/0006-3568(2001)051[0933:TEOTWA]2.0.CO;2.

### Alternatives in model type and specifications

The three antecedent effect models we present in the article are the subset of the potential models that we considered useful for our dataset. Our alternatives comprised both model types and model predictors. With respect to model types, the exploratory models developed in the literature can be subdivided in sliding windows, weighted means, regularization, and machine learning (van de Pol and Bailey 2019). In this study, we discarded the sliding windows, and machine learning models. Moreover, we also discarded particular types of weighted mean models (WMM), and of regularization models based on splines (Teller et al., 2016). With respect to model predictors, we discarded models including nonlinear predictors, more than one climatic predictor and their interaction, density dependent terms, and predictors directly related to plant physiology. We detail the reasons for these decisions below.

We discarded sliding windows because they are optimized to work in a maximum likelihood, rather than a Bayesian, framework, and because too complex for our small datasets. We also unsuccessfully tested a Bayesian model to substitute the maximum likelihood sliding window model. Sliding window models select the discrete windows of time – such as the window between May and July – with the best predictive ability. Sliding windows models are implemented by fitting one model for each potential climate window, select the best climate window via AIC model selection, and then test for the statistical significance of the selected climate window through randomization tests (van de Pol et al., 2016). Implementing this procedure in our Bayesian framework had three major issues. First, sliding windows are too computationally expensive in a Bayesian framework where performance is assessed with a leave-one-year-out cross-validation. Second, results cannot be directly compared to our three antecedent effect models, which essentially weigh the importance of each monthly anomaly. Third, the sliding windows models rely on significance tests via randomization simulations that do not make sense in a Bayesian framework.

To obviate the issues related to sliding window models, in our preliminary analyses we unsuccessfully tested a WMM designed to mimic the behaviour of a sliding window model. This WWM employed a generalized normal distribution whose scale parameter was fixed at 20. Such scale parameter imposes a rectangular shape to the generalized normal distribution, and thus a constant weight for the months located within one standard deviation of the mean. This model mimics a sliding window approach because it identifies the best window of climate predictors by estimating its center (through the mean) and width (through the standard deviation), and within this window each month has the same weight (e.g. Fig. 1a in van de Pol and Cockburn 2011). Unfortunately, this modified WWM using a generalized normal distribution had persistent convergence issues. We hypothesize this happened because convergence requires alternative climate windows to be adjacent.

We discarded machine learning models as not easily implemented in a Bayesian framework, and too data intensive for our relatively small datasets. Machine learning models tend to overfit data, so to minimize this problem they often rely k-fold or holdout cross-validation. Our datasets were so small that a common 10-fold cross-validation was often either not possible, or coincided with our leave-one-year-our cross-validation.

The WMMs we implement in our study are simplified with respect to what originally proposed by van de Pol and Cockburn (2011) to allow for model convergence. We fit our WMMs using a gaussian distribution. However, van de Pol and Cockburn (2011) had originally recommended fitting WMMs via the Weibull and generalized extreme value probability distributions. The advantage of these two distributions is the flexibility of their shape. However, we discarded these distributions because in our tests they consistently failed to converge. We hypothesize that these convergence issues might reflect low sample sizes, rather than a flaw in model specification.

We discarded the spline models proposed by Teller et al. (2016) because they were redundant with our three antecedent models. Specifically, the flexibility of spline models is intermediate between WMMs and Finnish Horseshoe Model (FHM). Given this redundancy, and given technical difficulties in implementing splines in a Bayesian framework, we discarded these models.

We discarded a set of potential alternative model predictors, many of which would have potentially led to overfitting given our low sample sizes. We discarded nonlinear predictors as we rarely saw evidence for nonlinear effects in the literature. We also discarded models including the interaction of temperature and precipitation. We did so because interactions amplify estimated uncertainty so much that sample sizes should be increased by an order of magnitude (Ch 16.; Gelman et al. 2020). We could not include density dependence, because our database was made entirely of density-independent matrix projection models. Finally, we chose to discard alternative climatic predictors such as growing degree days, and soil moisture, and standardized precipitation-evapotranspiration index. Our preliminary analyses with these predictors surprisingly showed a lower, rather than higher predictive ability than temperature or precipitation anomalies (data not shown).

### Models for data on survival, development, and reproduction

We modified the five models we fit on log(λ) data to fit survival, development, and reproduction data, which are non-normally distributed variables. These modifications are required for two reasons that are linked to the characteristics of the beta- (survival and development) and gamma (reproduction) distributions. First, survival and development represent transition probabilities bounded between zero and one, and reproduction a non-negative quantity bounded between 0 and positive infinity. We modelled these variables using, respectively, a logit and log link functions. Second, we adopted parameterizations that allowed us to model the mean of these variables. We illustrate our modifications using the CSM (Eq. 2) as an example. First, survival and development are rates that vary from 0 to 1, so we modeled these response variables using beta-regressions:

$$y_{ti} \sim \text{Beta}(a, b), \qquad \text{(S1a)}$$

$$a = \hat{y}_t \varphi \qquad \text{(S1b)}$$

$$b = (1 - \hat{y}_t)\varphi \qquad \text{(S1c)}$$

$$\text{logit}(\hat{y}_t) = \alpha + \beta \bar{x}_t \qquad \text{(S1d)}$$

$$\varphi \sim Gamma(1,1), \qquad \text{(S1e)}$$

where $\hat{y}_t$ refers to the mean prediction of either survival or development in year *t*, $\varphi$ represents the scale of the data, *a* and *b* are the shape parameters of the Beta distribution, and $y_{ti}$ is a single survival or development value observed in year *t* and spatial replicate *i*.

We modeled reproduction as gamma distributed, in this case using a log link that ensures non-negative predictions,

$$y_{ti} \sim \text{Gamma}\left(\kappa, \frac{\kappa}{\hat{y}_t}\right), \qquad \text{(S2a)}$$

$$\log(\hat{y}_t) = \alpha + \beta \bar{x}_t. \qquad \text{(S2b)}$$

$$\kappa \sim Gamma(0.01, 0.01). \qquad \text{(S2c)}$$

In Eq. S2, $\hat{y}_t$ refers to the mean prediction for reproduction values in year *t*, $\kappa$ is the shape parameter of the gamma distribution, and $\kappa/\hat{y}_t$ is its scale calculated through the maximum likelihood estimator, and $y_{ti}$ refers to a single reproduction value observed in year *t* and spatial replicate *i*.

### Antecedent effect model infographic

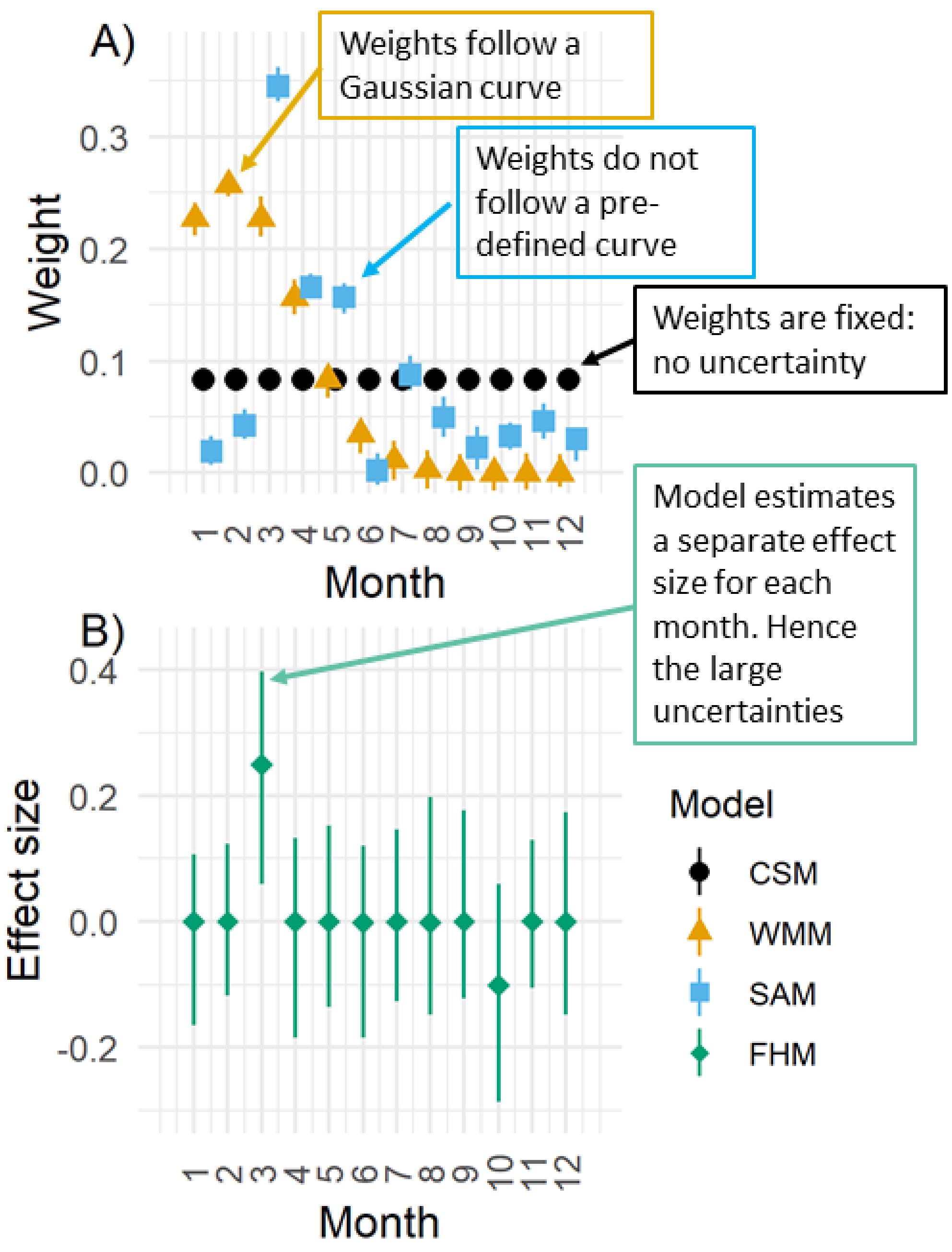

Figure S1. Antecedent effect models used to estimate the effect of monthly climate anomalies on population dynamics. These plots represent hypothetical Bayesian estimates of the A) weight and B) effect sizes of 12 monthly climate anomalies observed before a demographic outcome (*e.g.*, change in survival). In both panels, the symbols represent estimate means, and the extremes of the error bars represent the 95% credible intervals. In panel A), the effect size of climate depends on the relative weights of monthly anomalies. The Climate Summary Model (CSM) is a linear regression where every monthly anomaly has the same weight; the climatic predictor in this regression is thus given by the mean of 12 monthly anomalies. Hence, monthly weights are identical, and no error bars are shown. In the Weighted Mean Model (WMM), monthly weights follow a Gaussian curve; in the Stochastic Antecedent Model (SAM) the monthly weights are determined by a Dirichlet distribution. Finally, panel B) shows how the Finnish Horseshoe Model (FHM) tends to produce effect sizes centered around zero – except for the few effect sizes, two in this hypothetical case, for which there is strong evidence of a positive and negative effect.

### Prior predictive checks

To compare the predictive ability of our five models, we performed simulations to identify weakly informative priors. These simulations are called "prior predictive checks", which represent the process of simulating data from prior distributions and assessing the results, sometimes comparing them to the observed data (Gabry et al., 2019). We chose the priors that produced data not greatly exceeding the lower and upper extremes of the observed data (Gabry et al., 2019). We call the priors chosen through this process "weakly informative" priors, following Gabry et al. (2019). These not only facilitate direct comparison of model predictive ability, but they also facilitate model convergence. To show how we performed the prior predictive checks, we use the simple linear regression model (Eq. 2 in the main text) as an example. This model is defined as:

$$y_{ti} \sim Normal(\hat{y}, \sigma), \qquad \text{(S3a)}$$

$$\hat{y}_t = \alpha + \beta \bar{x}_t, \qquad \text{(S3b)}$$

$$\alpha \sim Normal(0{,}0.5), \qquad \text{(S3c)}$$

$$\beta \sim Normal(0{,}1), \qquad \text{(S3d)}$$

$$\sigma \sim Gamma(0.01{,}0.01) \qquad \text{(S3e).}$$

Here, the priors were in Eq. S1c-e, and $\bar{x}_t$ refer to nine years of randomly generated, normally distributed, climate predictors. We used the priors in Eq. S1c-e to simulate 60,000 values of $y_{ti}$ as generated by the climate summary model (CSM, Eq. 2). We also ran simulations for the remaining four models: the null model (NM, Eq. 1), weighted mean model (WMM, Eq. 3), stochastic antecedent model (SAM, Eq. 4), and Finnish horseshoe model (FHM, Eq. 5). The CSM, WMM, and SAM shared the same priors shown in Eq. S3 for $\alpha$, $\beta$, and $\sigma$; the NM shared the priors shown in Eq. S3 for $\alpha$, and $\sigma$. Note that in Eq. S3c, the prior for $\alpha$ has a lower standard deviation than the prior for $\beta$. This difference is justified because $\alpha$ had a larger effect than $\beta$ on the variance of $\hat{y}_t$ values (data not shown). The prior for the FHM differed because instead of a single $\beta$ parameter, it was comprised of 12 $\beta_k$ parameters (Eq. 5b). Reproducing Eq. 5 in the main text, $\beta_k$ is defined as:

$$\beta_k \sim Normal(0, \tau\tilde{\theta}_k), \qquad \text{(S4a)}$$

$$\tilde{\theta}_k = \frac{c\theta_k}{\sqrt{c^2 + \tau^2\theta_k^2}}, \qquad \text{(S4b)}$$

$$\theta_k \sim HalfCauchy(0, 0.1), \qquad \text{(S4c)}$$

$$c^2 \sim InvGamma\left(\frac{v}{2}, \frac{v}{2}s^2\right), \qquad \text{(S4d)}$$

$$\tau \sim HalfCauchy(0, \tau_0), \qquad \text{(S4e)}$$

$$\tau_0 = \frac{m_0}{M - m_0}\frac{\sigma}{\sqrt{N}}. \qquad \text{(S4f)}$$

Here, the standard deviation of each $\beta_k$ value is one order of magnitude smaller than the $\beta$ parameter in Eq. S3d. A *HalfCauchy* distribution with a standard deviation of 1 would increase the range of $\hat{y}_t$ much beyond the observed data (data not shown). We compared the distributions of data simulated via the priors described above to the 982 log(λ) values linked to our 34 study species. The priors in Eq. S3c-e simulate a large proportion of values falling within the range of observed data (Figure S1).

We used the same simulation procedure to find informative priors for the beta models (Eq. S1a-e, used to fit survival and growth data) and gamma models (Eq. 2a-c, used to fit reproduction data). In the beta model, these priors

$$\alpha \sim Normal(0,1), \qquad \text{(S5a)}$$

$$\beta \sim Normal(0,1), \qquad \text{(S5b)}$$

$$\varphi \sim Gamma(1,1), \qquad \text{(S5c)}$$

simulated a range of values similar to the 1,964 observed survival and progression data points (Figure S2). Here, the prior for the dispersion parameter $\varphi$ produces lower dispersion when compared to the prior of $\sigma$ (Eq. S5c versus Eq. S3e). Low dispersion is needed in beta models to avoid generating $\hat{y}_t$ values concentrated on 0 and 1 (Hobbs & Hooten, 2015). Therefore, we decreased the dispersion in beta models by defining the prior for the dispersion parameter as $\varphi \sim Gamma(1,1)$.

In gamma models, these priors

$$\alpha \sim Normal(0,0.5), \qquad \text{(Eq. S6a)}$$

$$\beta \sim Normal(0,1), \qquad \text{(Eq. S6b)}$$

$$\kappa \sim Gamma(0.01,0.01). \qquad \text{(Eq. S6c)}$$

simulated realistic values when compared to the 982 data points on reproduction (Figure S3). We carried out all simulations in R, which facilitated simulating truncated distributions through the package truncdist (Novomestky & Nadarajah, 2016). We used a truncated distribution to simulate the half-cauchy distribution associated with the Finnish Horseshoe model (Eq. 5d and 5f).

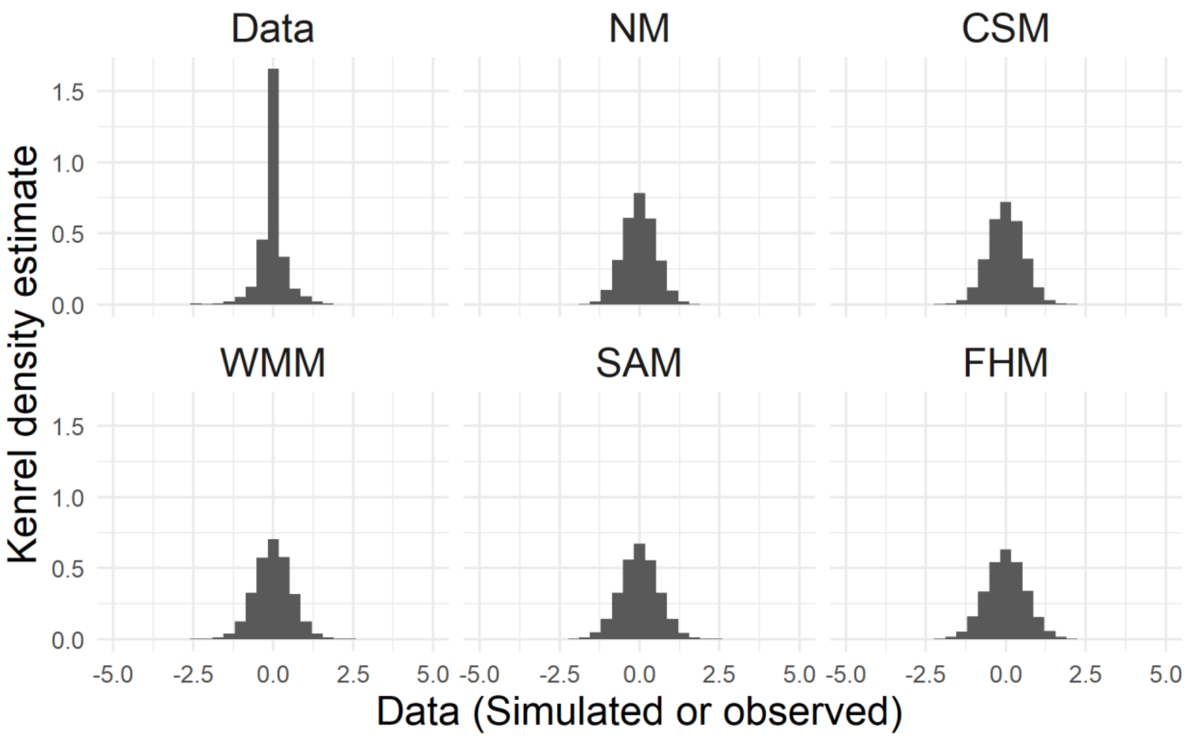

Figure S2. Kernel density estimate of the 982 observed population growth rate, log($\lambda$), values (upper left panel), and of the 60,000 values simulated using priors in Eq. S3-4 (remaining panels). The acronyms used as panel titles refer to null models (NM), climate summary models (CSM), weighted mean models (WMM), stochastic antecedent models (SAM), and Finnish horseshoe Model (FHM).

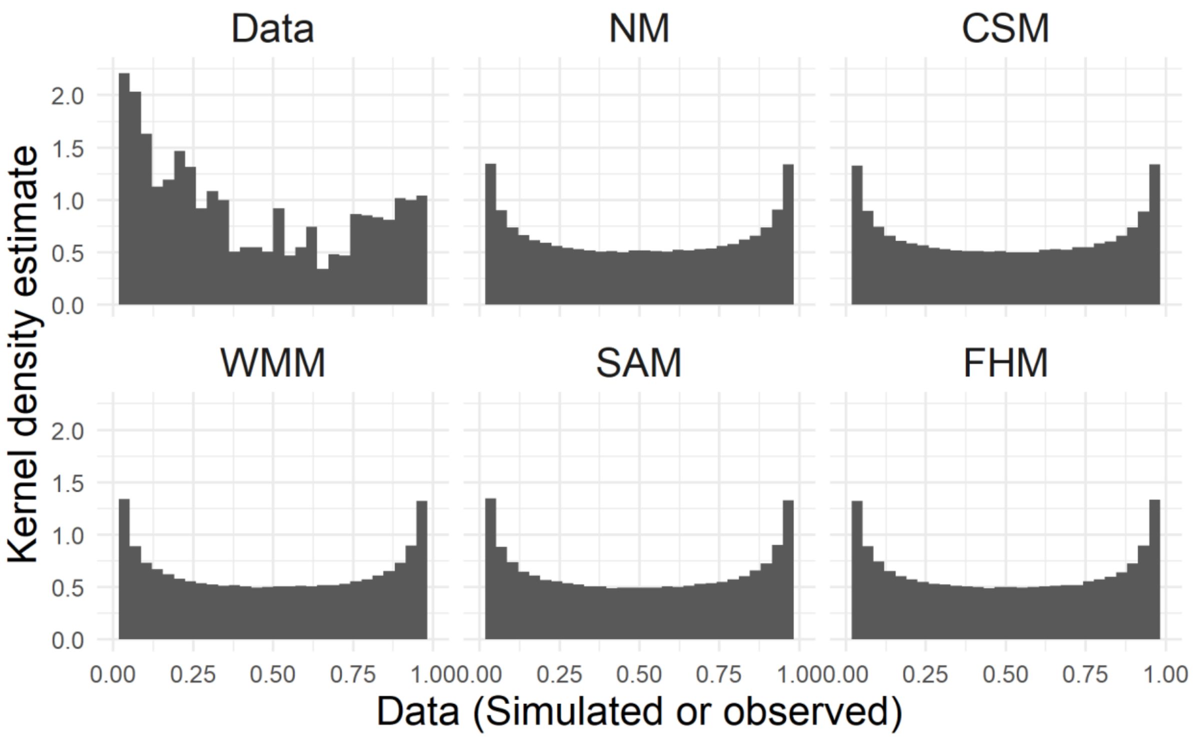

Figure S3. Kernel density estimates of the 1964 observed survival and growth values (upper left panel), and of the 60,000 values simulated using priors in Eq. S5 (all other panels). The acronyms used as panel titles refer to null models (NM), climate summary models (CSM), weighted mean models (WMM), stochastic antecedent models (SAM), and Finnish horseshoe Model (FHM).

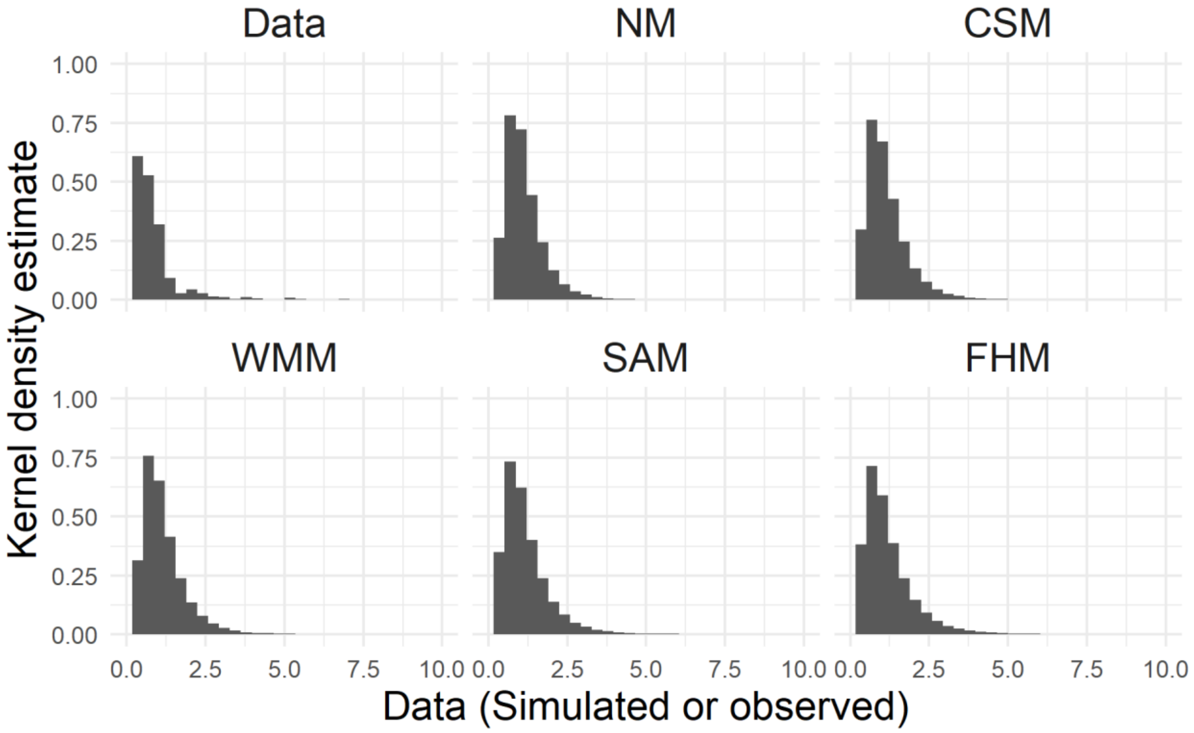

Figure S4. Kernel density estimates of the 982 observed reproduction values (upper left panel), and of the 60,000 values simulated using priors in Eq. S6 (all other panels). The acronyms used as panel titles refer to null models (Null), climate summary models (CSM), weighted mean models (WMM), stochastic antecedent models (SAM), and Finnish horseshoe Model (FHM).

### Model convergence and model checks

We chose four separate diagnostics to identify potential issues in model fitting. First, divergent transitions are a specific diagnostic for Hamiltonian Monte Carlo samplers. These samplers approximate continuous functions using discrete steps. Divergent transitions occur when the discrete steps employed by the sampler are too large to approximate the underlying continuous function (Stan Development Team, n.d.). We flagged models with more than 10 divergent transitions because to diagnose a problem with the sampler, multiple divergent transitions need to occur in the same location of parameter space. Second, the Gelman and Rubin convergence diagnostic measures whether the variance of parameter values within single MCMC chains is comparable to the variance among chains. Larger variance among chains indicate convergence issues (Gelman & Rubin, 1992). Third, we considered the Monte Carlo standard error of a parameter as high when the ratio of Monte Carlo standard error to the standard deviation of the posterior was above 0.1. Fourth, the effective sample size decreases with an increase in the autocorrelations of MCMC chains (Gelman et al., 2013). We considered the effective sample size of a parameter low whenever it was below 10% of the total 6,000 posterior samples for our models.

An alternative way to identify convergence issues is to plot the $\hat{R}$ versus the effective sample size of each parameter. Parameters with convergence issues tend to have high $\hat{R}$ and low effective sample size, an issue easily identified in bivariate plots. We produced these diagnostic plots for each model type and climatic predictor.

We further tested that our models represented the data well by performing model Bayesian checks. We checked our models by calculating the Bayesian p-values of posterior predictive checks. Bayesian p-values are more conservative, and should be preferred to, posterior predictive checks based on a loss functions such as sum of squared residuals (Conn et al., 2018). To perform a sampled posterior predictive check, we extracted one random sample from the joint posterior of each model to produce 1,000 simulated datasets. We obtained Bayesian p-values by calculating the proportion of times the sum of squared residuals (SSR) of the simulated data exceeded that of observed data. A proportion between 0.025 and 0.975 suggests the model reproduces the observed data well.

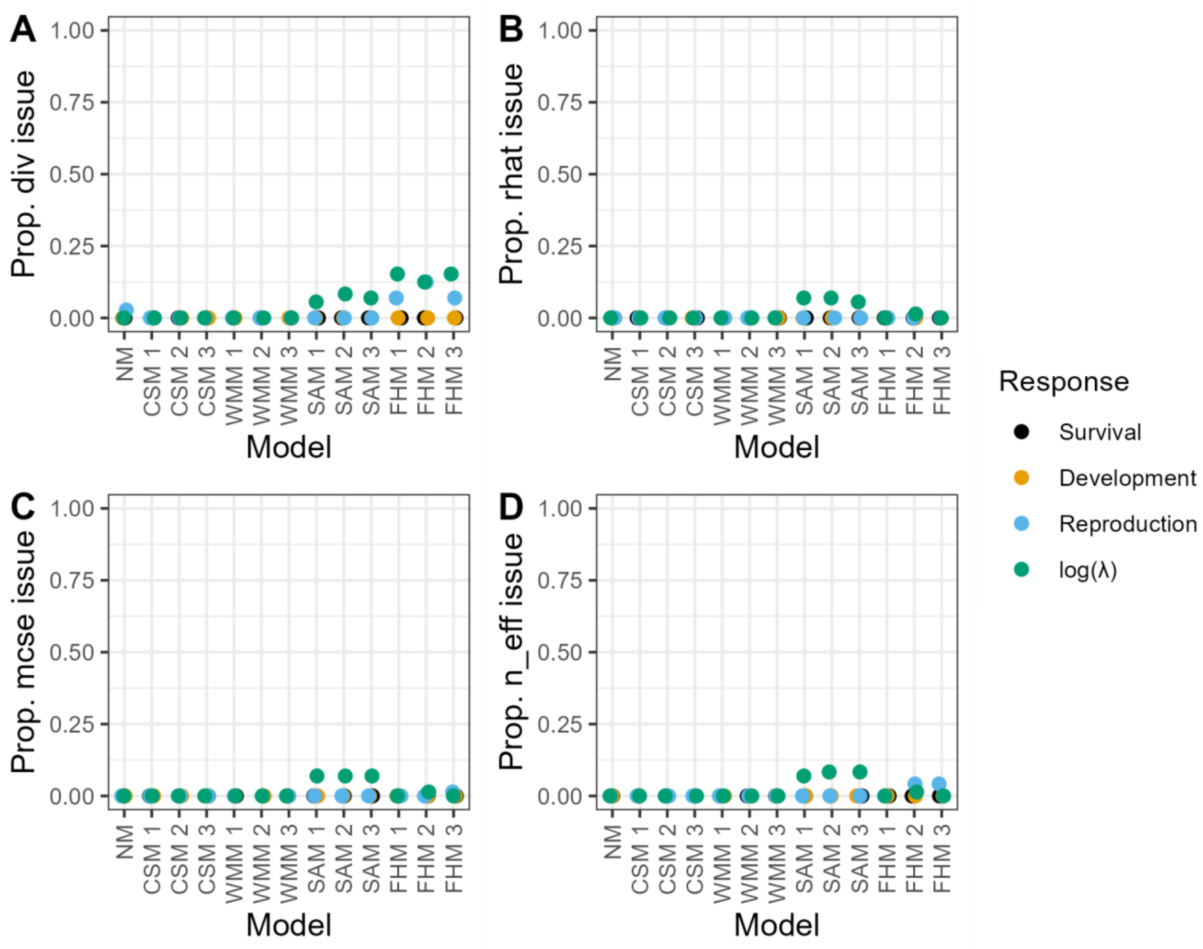

Figure S5. Few models showed problems with convergence. Proportion of models that had at least (A) one divergent transition, (B) one model parameter whose R̂ exceeded 1.1, (C) one model parameter whose MSCE was below 10%, and (D) one model parameter whose effective sample number was less than 600. The labels of the x-axis are acronyms that refer to model type: the Climate Summary Model (CSM), Weighted Mean Model (WMM), Stochastic Antecedent Model (SAM), and Finnish Horseshoe Model (FHM). The numbers refer to the year of climate data used.

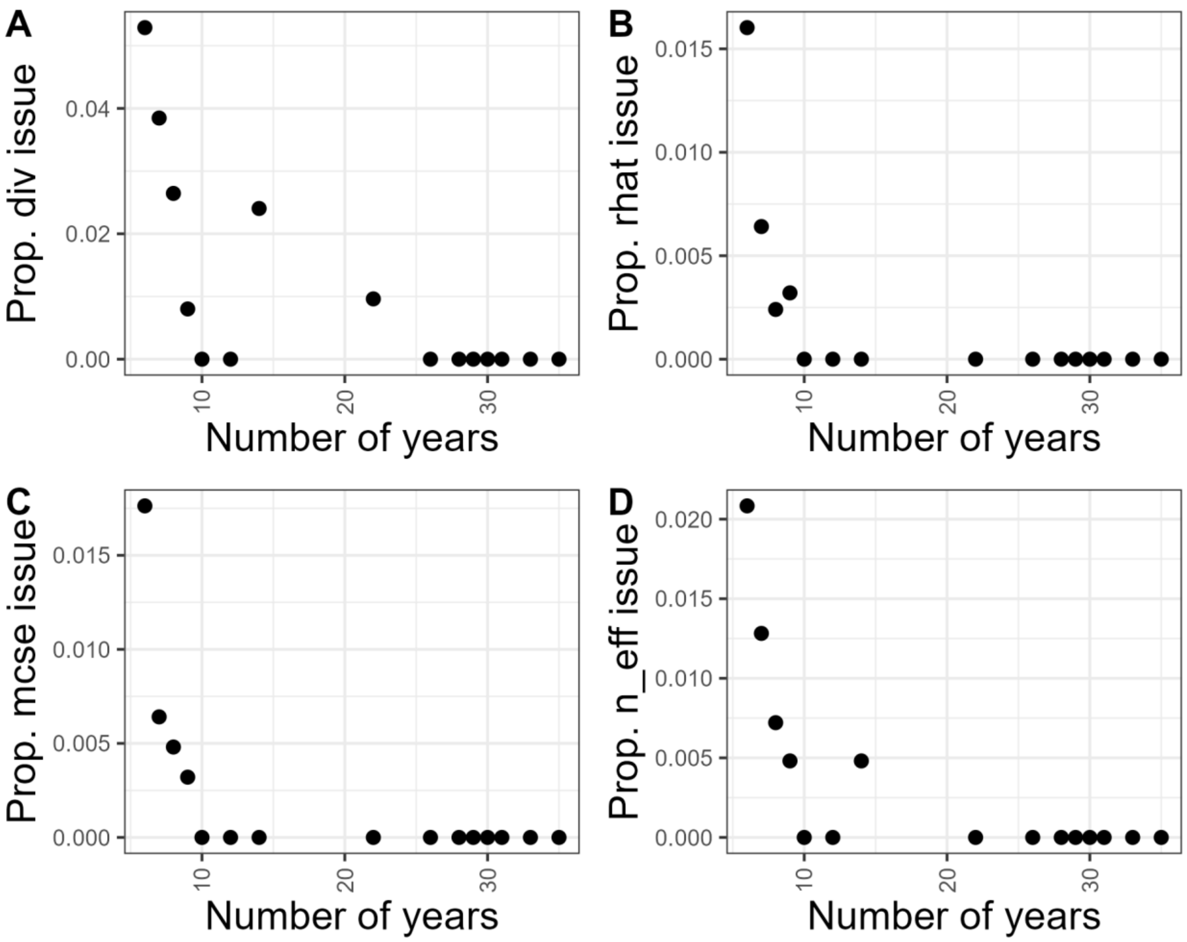

Figure S6. Convergence issues were concentrated in datasets with less than 20 years of data. Using the number of years of data available for each of our 36 studies on the x-axis, scatter plots showing the average proportion of models that had at least (A) one divergent transition, (B) one model parameter whose $\hat{R}$ exceeded 1.1, (C) one model parameter whose MSCE was below 10%, and (D) one model parameter whose effective sample number was less than 600.

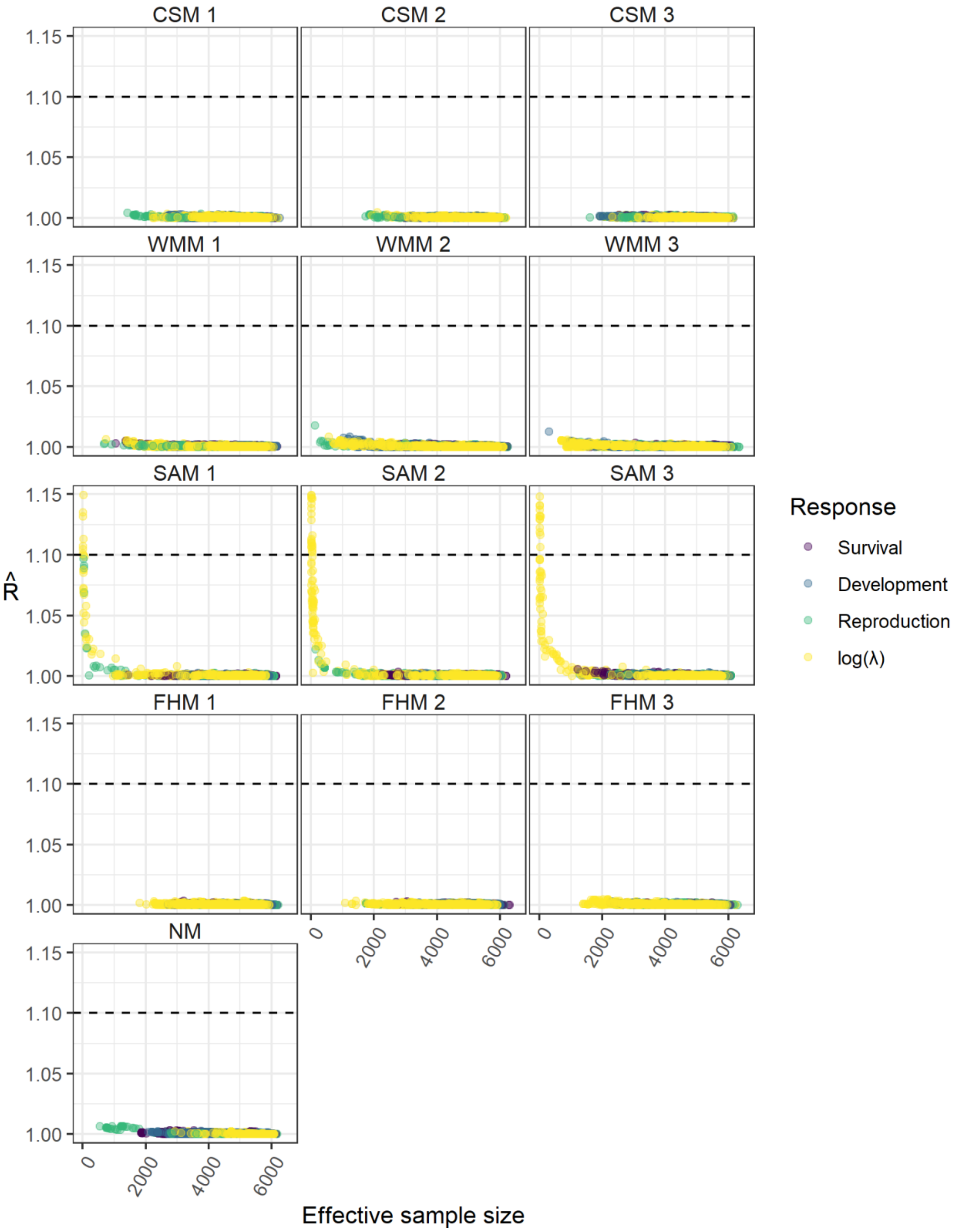

Figure S7. Gelman Rubin convergence factor ($\hat{R}$) vs. effective sample size for parameters of the 12 models fit using air temperature as a predictor. The title of each panel refers to model type, with the acronyms referring to the Climate Summary Model (CSM), Weighted

Mean Model (WMM), Stochastic Antecedent Model (SAM), and Finnish Horseshoe Model (FHM). The numbers refer to the year of climate data used.

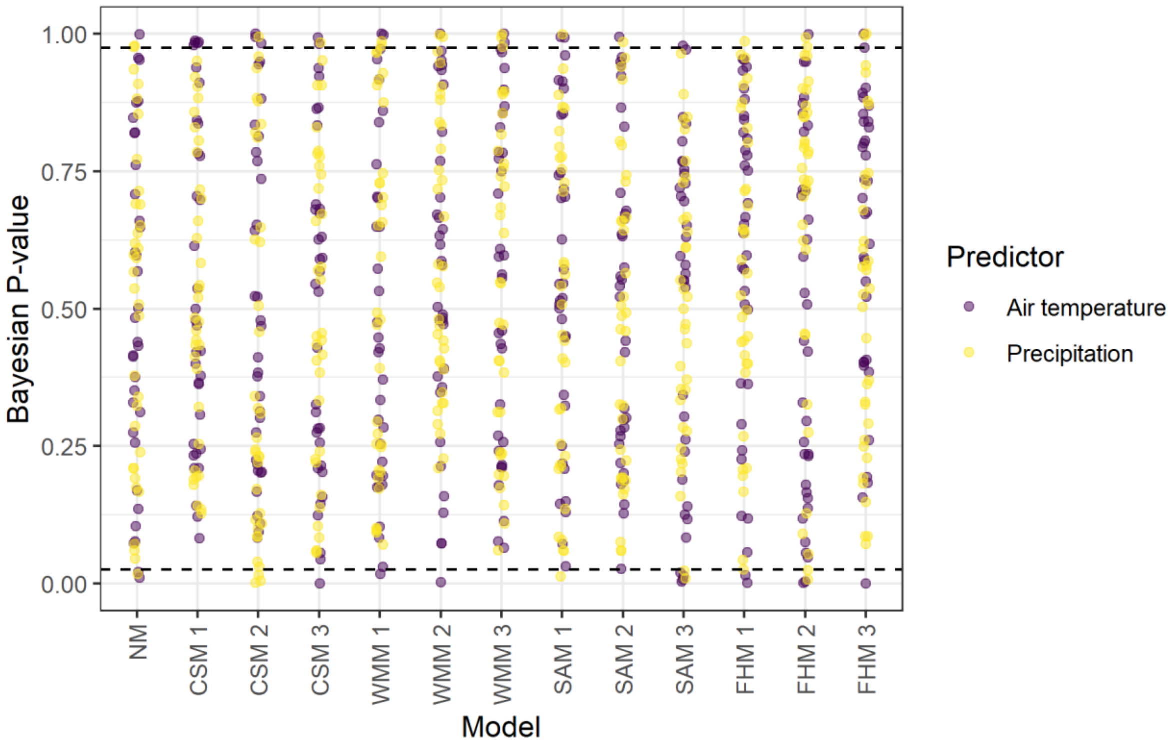

Figure S8. Successful posterior predictive checks. Bayesian p-values associated with sampled posterior predictive checks on the 13 models fit using log($\lambda$) as a response variable, and air temperature (violet) and precipitation (yellow) as predictors. Upper and lower dotted line represent the 97.5% and 2.5% percentiles, respectively. The labels of the x-axis are acronyms referring to model types: the Climate Summary Model (CSM), Weighted Mean Model (WMM), Stochastic Antecedent Model (SAM), and Finnish Horseshoe Model (FHM). The numbers refer to the year of climate data used.

### Results figures

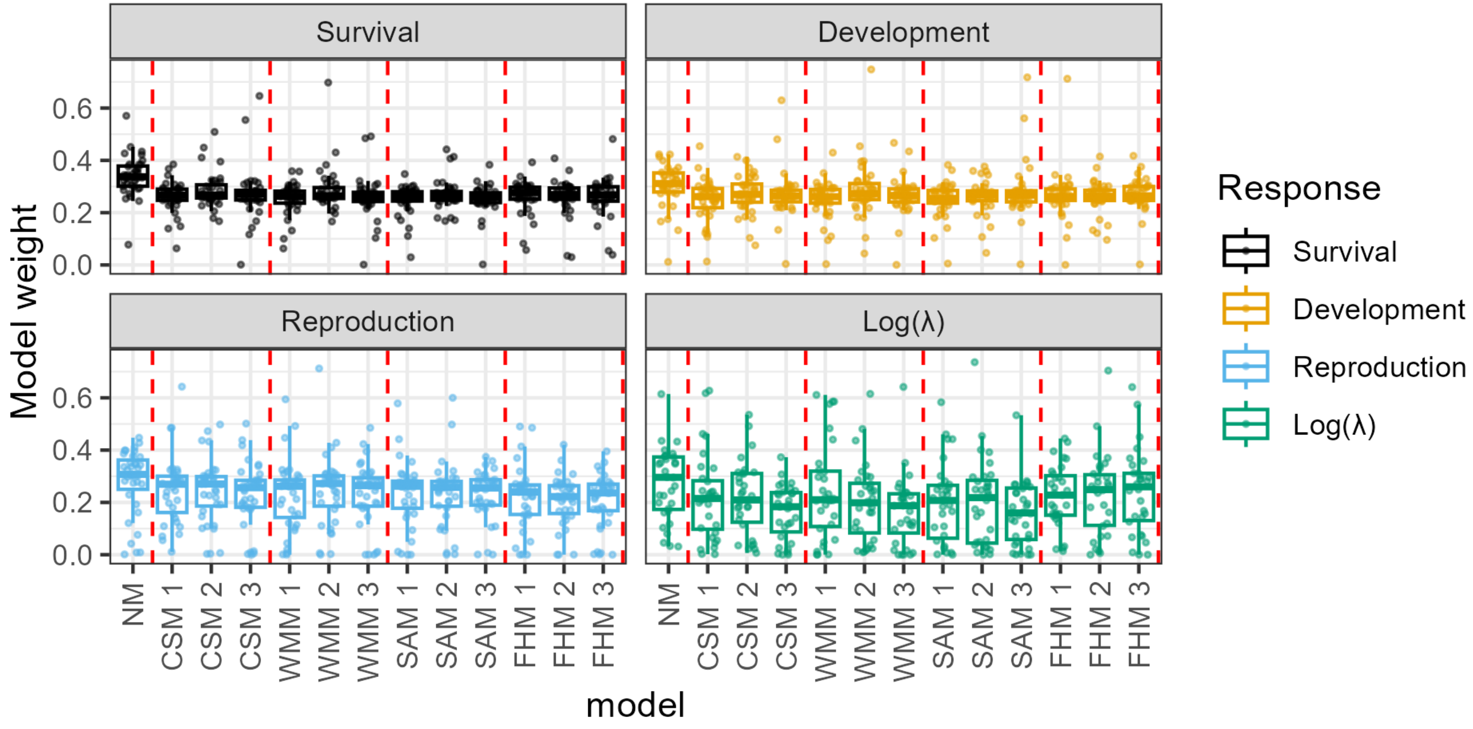

Figure S10. Climate models do not provide sensible advantages when predicting vital rates using precipitation as predictor. Box and whisker plots and point clouds showing the arcsine transformed model weights for our 34 species, across five model types, three periods (1: year *t*, 2: year *t*-1, 3: year *t*-2), and four response variables. The middle line of the boxplots shows the median, the upper and lower hinges delimit the first and third quartile, and the whiskers extend 1.5 times beyond the first and third quartile. Each point refers to a single model fit.

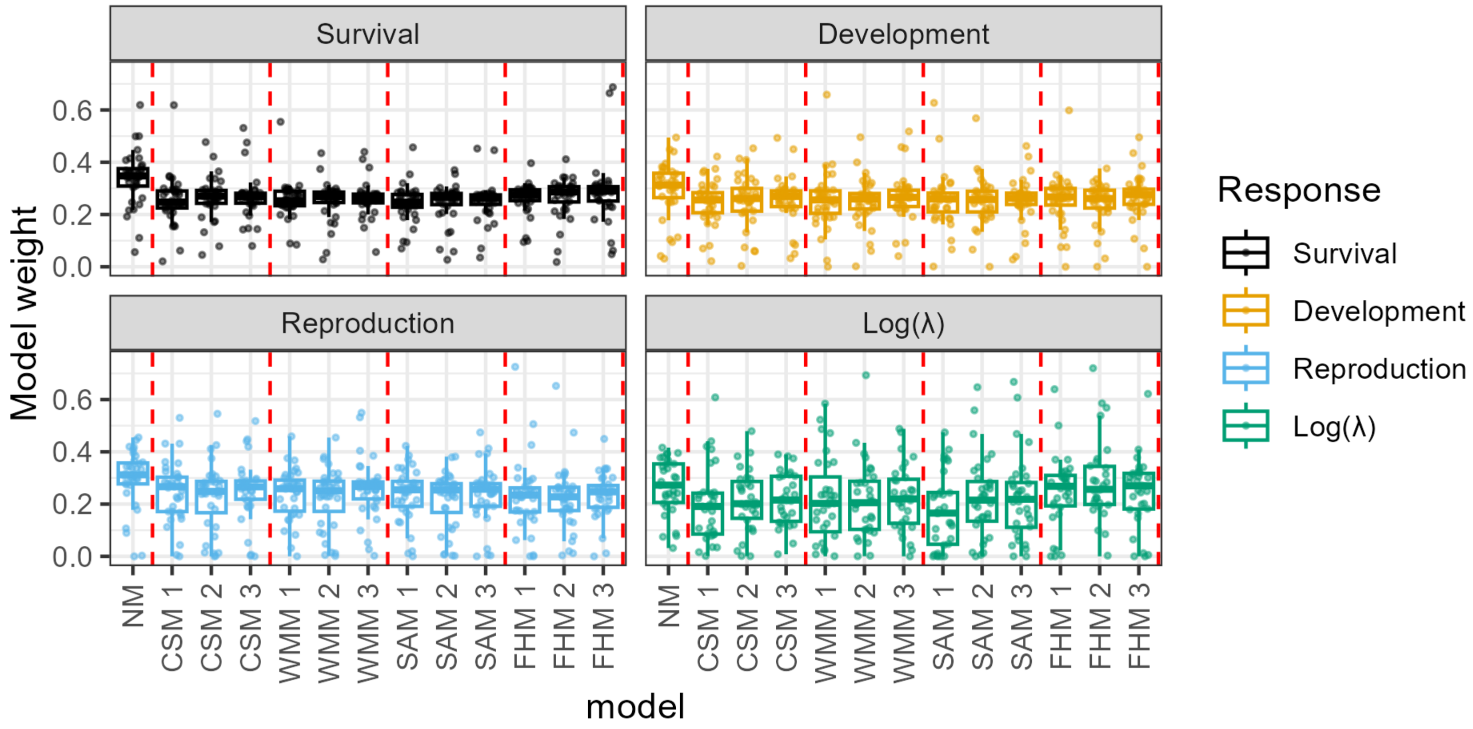

Figure S11. Climate models do not provide sensible advantages when predicting vital rates using temperature as predictor. Box and whisker plots and point clouds showing the arcsine transformed model weights for our 34 species, across two climatic predictors, five model types, three periods (1: year *t*, 2: year *t*-1, 3: year *t*-2), and four response variables. The middle line of the boxplots shows the median, the upper and lower hinges delimit the first and third quartile, and the whiskers extend 1.5 times beyond the first and third quartile. Each point refers to a single model fit.

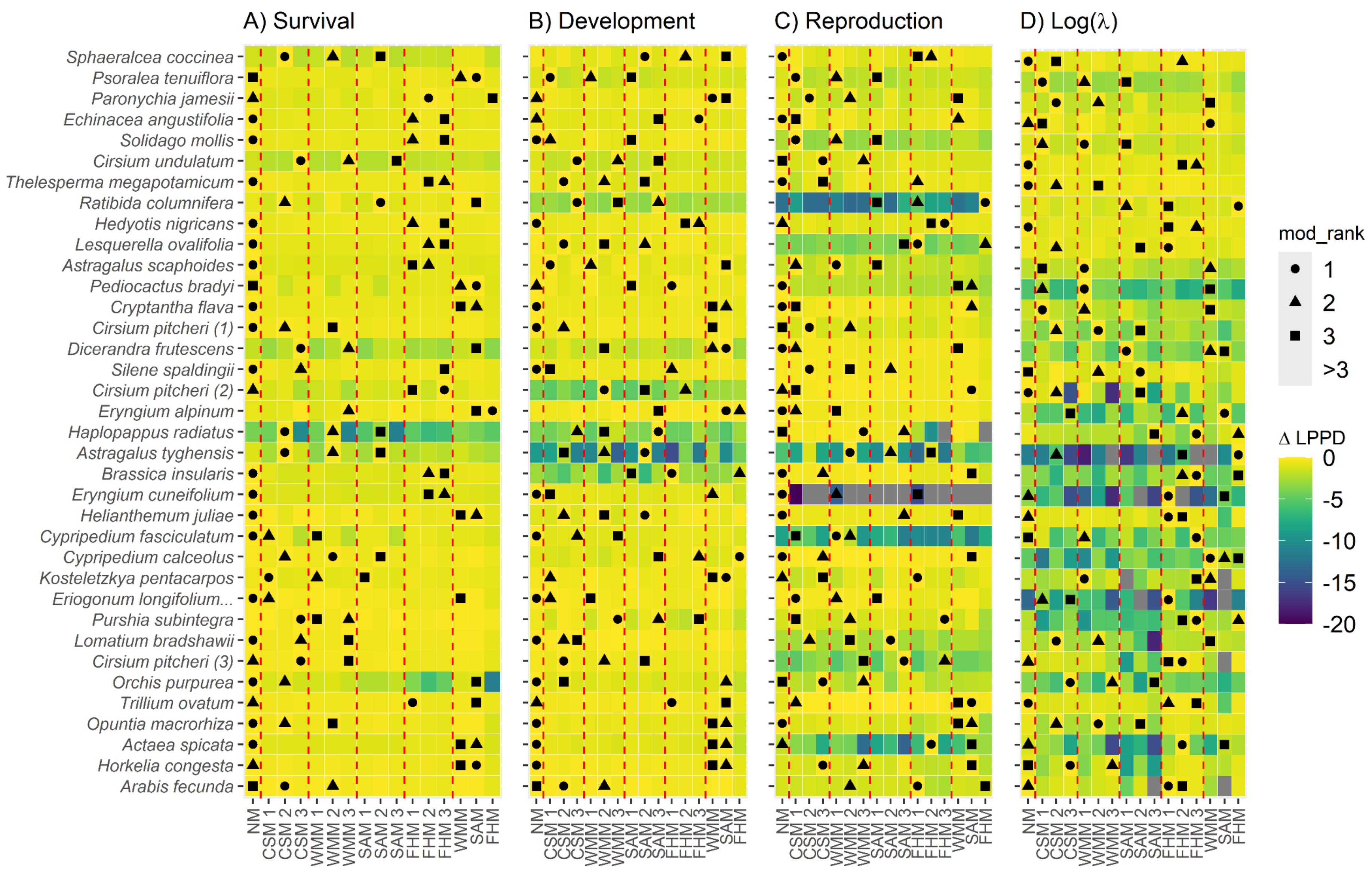
A) Survival
B) Development
C) Reproduction
D) Log(λ)
Sphaeralcea coccinea
Psoralea tenuiflora
Paronychia jamesii
Echinacea angustifolia
Solidago mollis
Cirsium undulatum
Thelesperma megapotamicum
Ratibida columnifera
Hedyotis nigricans
Lesquerella ovalifolia
Astragalus scaphoides
Pediocactus bradyi
Cryptantha flava
Cirsium pitcheri (1)
Dicerandra frutescens
Silene spaldingii
Cirsium pitcheri (2)
Eryngium alpinum
Haplopappus radiatus
Astragalus tyghensis
Brassica insularis
Eryngium cuneifolium
Helianthemum juliae
Cypripedium fasciculatum
Cypripedium calceolus
Kosteletzkya pentacarpos
Eriogonum longifolium...
Purshia subintegra
Lomatium bradshawii
Cirsium pitcheri (3)
Orchis purpurea
Trillium ovatum
Opuntia macrorhiza
Actaea spicata
Horkelia congesta
Arabis fecunda
NM
CSM 1
CSM 2
CSM 3
WMM 1
WMM 2
WMM 3
SAM 1
SAM 2
SAM 3
FHM 1
FHM 2
FHM 3
WMM
SAM
FHM
mod_rank
1
2
3
>3
Δ LPPD
0
-5
-10
-15
-20

Figure S12. Tile plot of predictive power based on differences in log pointwise predictive density (ΔLPPD) for models using precipitation as predictor. Tiles with yellow colors denote models with better predictive ability; the best models have a ΔLPPD of 0. Symbols denote best (circles), second best (triangle), and third best (square) model. Each panel refers to a separate response variable: A) survival, B) development, C) reproduction, and D) log($\lambda$). Rows refer to each dataset, and columns to each model. We compare five types of models (eqs. 1-5). We fit these models using the climate observations from three years prior to a demographic response. Species are organized by the temporal replication of the dataset, with the longest datasets being on top.

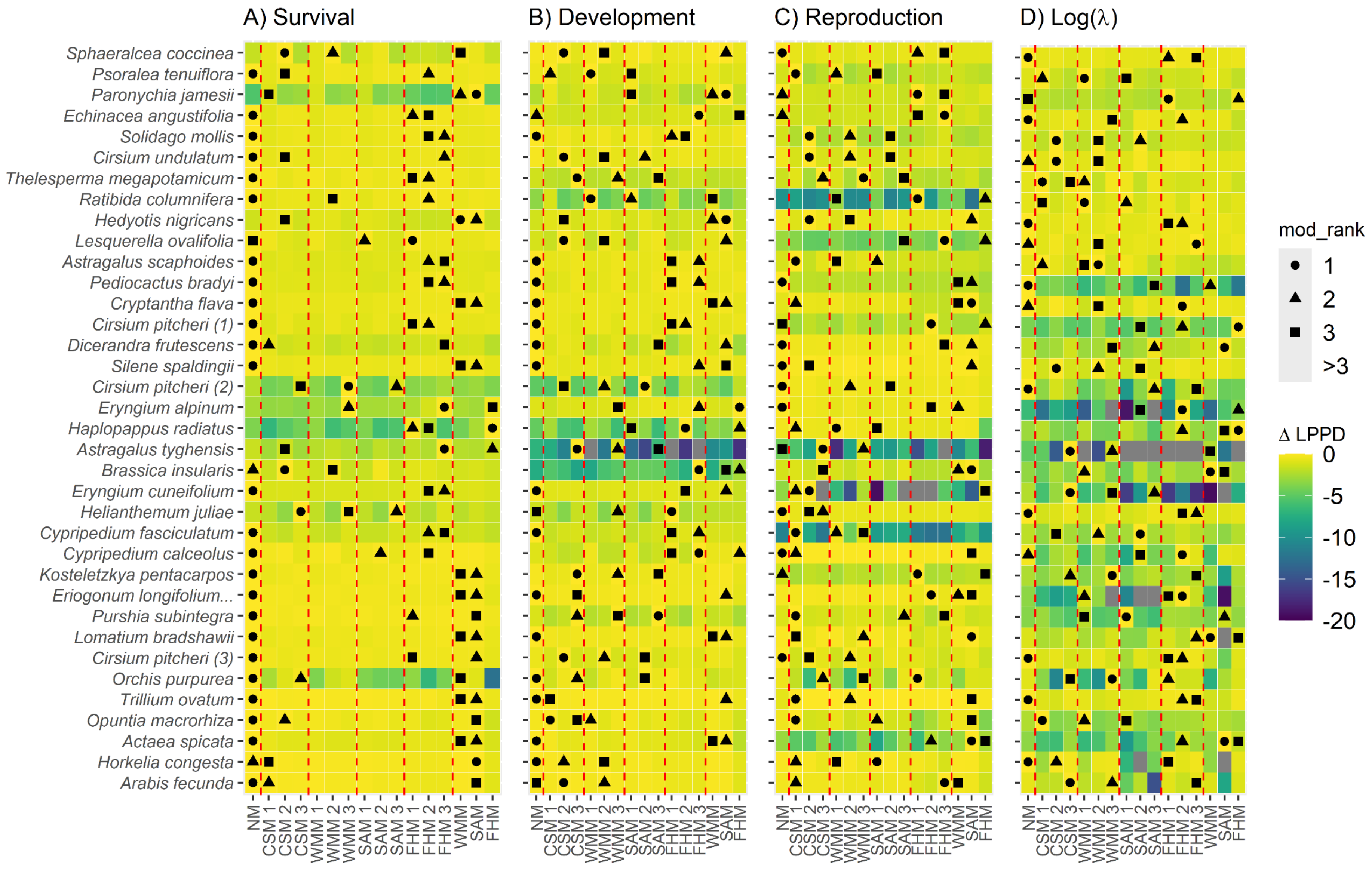

A) Survival
B) Development
C) Reproduction
D) Log(λ)
Sphaeralcea coccinea
Psoralea tenuiflora
Paronychia jamesii
Echinacea angustifolia
Solidago mollis
Cirsium undulatum
Thelesperma megapotamicum
Ratibida columnifera
Hedyotis nigricans
Lesquerella ovalifolia
Astragalus scaphoides
Pediocactus bradyi
Cryptantha flava
Cirsium pitcheri (1)
Dicerandra frutescens
Silene spaldingii
Cirsium pitcheri (2)
Eryngium alpinum
Haplopappus radiatus
Astragalus tyghensis
Brassica insularis
Eryngium cuneifolium
Helianthemum juliae
Cypripedium fasciculatum
Cypripedium calceolus
Kosteletzkya pentacarpos
Eriogonum longifolium...
Purshia subintegra
Lomatium bradshawii
Cirsium pitcheri (3)
Orchis purpurea
Trillium ovatum
Opuntia macrorhiza
Actaea spicata
Horkelia congesta
Arabis fecunda
NM
CSM 1
CSM 2
CSM 3
WMM 1
WMM 2
WMM 3
SAM 1
SAM 2
SAM 3
FHM 1
FHM 2
FHM 3
WMM
SAM
FHM
mod_rank
1
2
3
>3
Δ LPPD
0
-5
-10
-15
-20

Figure S13. Tile plot of predictive power based on differences in log pointwise predictive density (ΔLPPD) for models using temperature as predictor. Tiles with yellow colors denote models with better predictive ability; the best models have a ΔLPPD of 0. Symbols denote best (circles), second best (triangle), and third best (square) model. Each panel refers to a separate response variable: A) survival, B) development, C) reproduction, and D) log($\lambda$). Rows refer to each dataset, and columns to each model. We compare five types of models (eqs. 1-5). We fit these models using the climate observations from three years prior to a demographic response. Species are organized by the temporal replication of the dataset, with the longest datasets being on top.

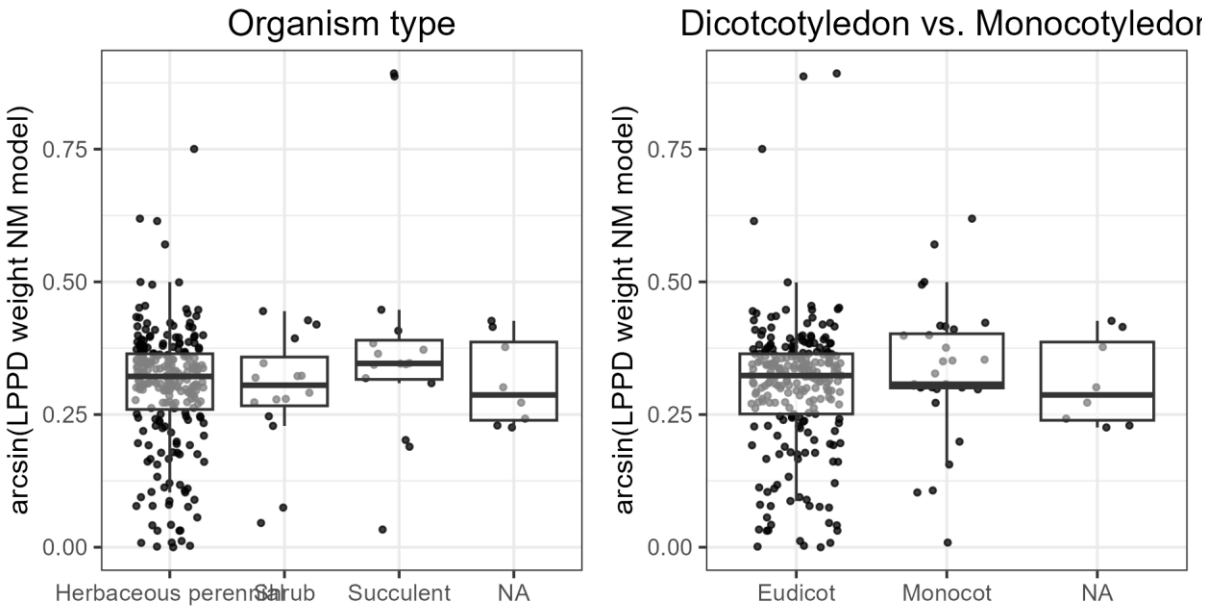

Figure S14. Plant functional types do not affect the predictive performance of climate models. Box and whisker plots and point clouds showing the arcsine transformed model weights of the null (NM) models by organism type (left panel), and plant group (Monocotyledons or Dicotyledons). Lower values reflect a higher predictive ability of the models including a climatic predictor. The middle line of the boxplots shows the median, the upper and lower hinges delimit the first and third quartile, and the whiskers extend 1.5 times beyond the first and third quartile.

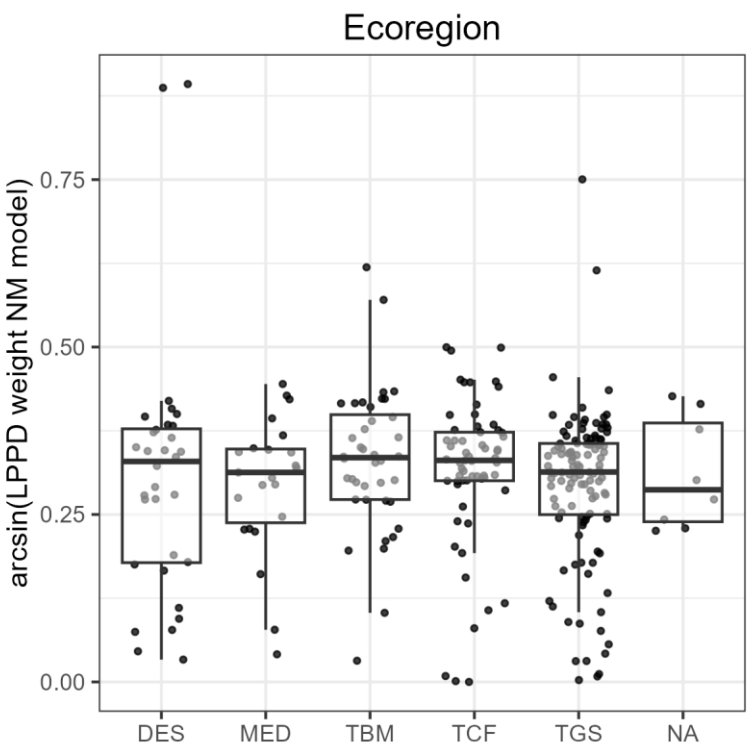

Figure S15. Ecoregion does not affect the predictive performance of climate models. Box and whisker plots and point clouds showing the arcsine transformed model weights of the null (NM) models by Ecoregion. Codes stand for: DES (Deserts And Xeric Shrublands), MED (Mediterranean Forests, Woodlands And Scrubs), TBM (Temperate Broadleaf And Mixed Forests), TCF (Temperate Grasslands, Savannas, And Shrublands), TGS (Temperate Coniferous Forests). Lower values reflect a higher predictive ability of the models including a climatic predictor. The middle line of the boxplots shows the median, the upper and lower hinges delimit the first and third quartile, and the whiskers extend 1.5 times beyond the first and third quartile.

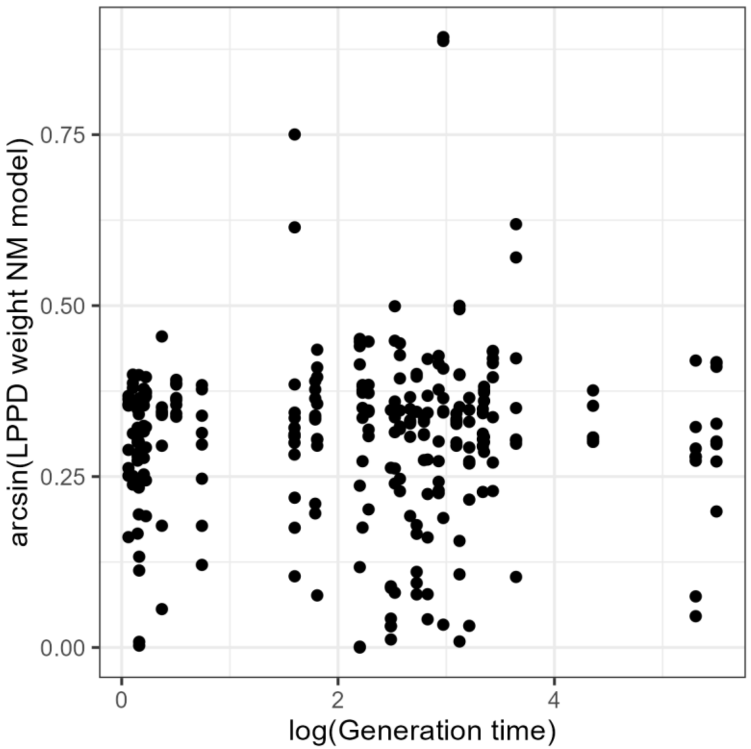

Figure S16. Generation time does not affect the predictive performance of climate models. Bivariate plot relating the arcsine transformed model weights of the null (NM) models with the natural logarithm of the generation time of the species models refer to. In this plot, a positive correlation between the two variables would imply climate tends to predict the demography of long-lived species better than null (NM) models.

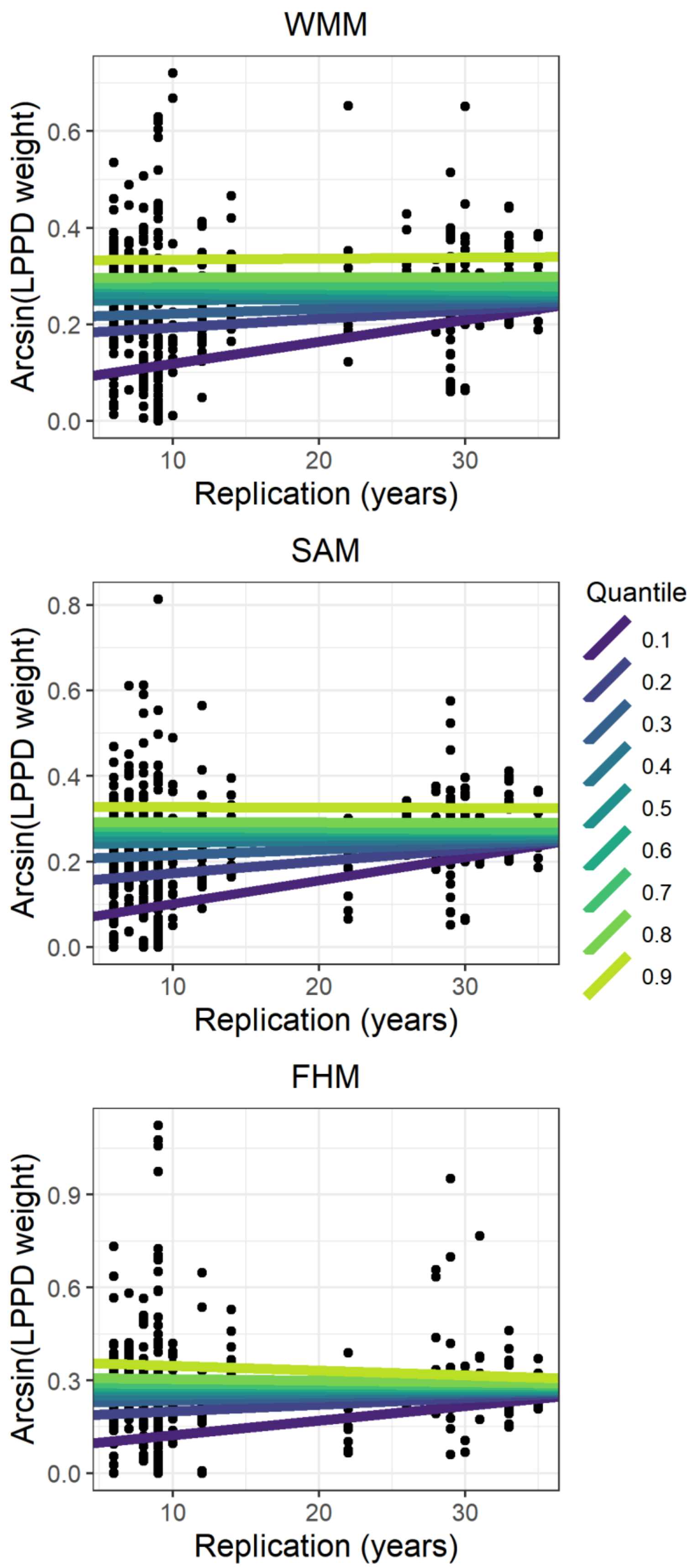

Figure S17. More years of data increase the predictive ability of the worst model in antecedent models. Scatterplots of the arcsine transformed model weights versus the years available to each dataset. Panels show the three types of antecedent effect models. The acronyms refer to the Weighted Mean Model (WMM), Stochastic Antecedent Model (SAM), and Finnish Horseshoe Model (FHM). Colored lines show quantile regression average predictions for nine quantiles, going from the 0.1 to the 0.9 quantile in increments of 0.1.

### Sensitivity analysis

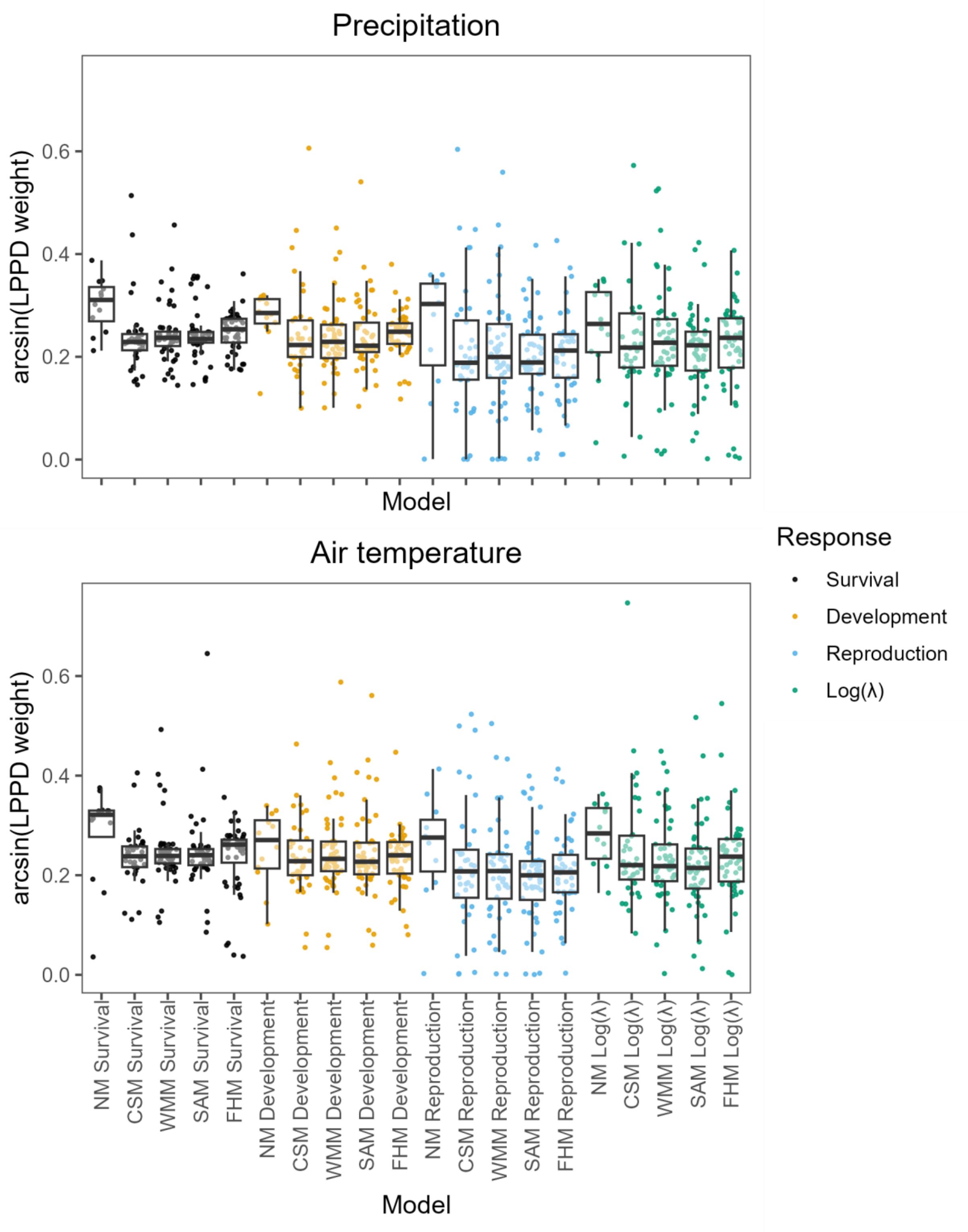

Figure S18. Null models have the best predictive ability in datasets with at least 20 years of data. Box and whisker plots and point clouds showing the arcsine transformed model weights for 12 species, across five model types, and four response variables. The middle line of the boxplots shows the median, the upper and lower hinges delimit the first and third quartile, and the whiskers extend 1.5 times beyond the first and third quartile. Each point refers to a single model fit.

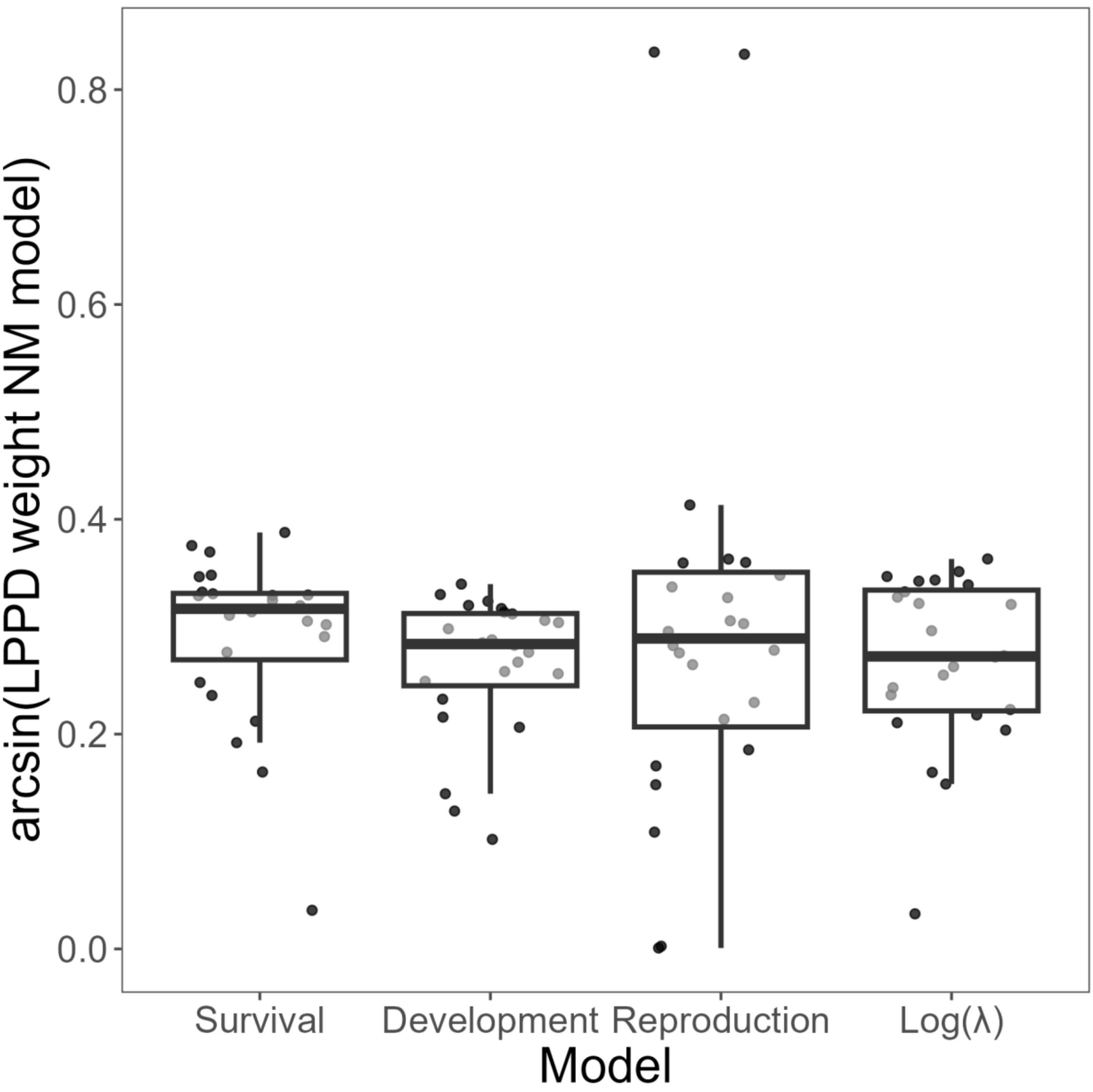

Figure S19. Climate has a similar predictive power across response variables in datasets with at least 20 years of data. Box and whisker plots and point clouds showing the arcsine transformed model weights of the null models based on the type of response variable. In this plot, lower values reflect a higher predictive ability of the models including a climatic predictor. The middle line of the boxplots shows the median, the upper and lower hinges delimit the first and third quartile, and the whiskers extend 1.5 times beyond the first and third quartile.

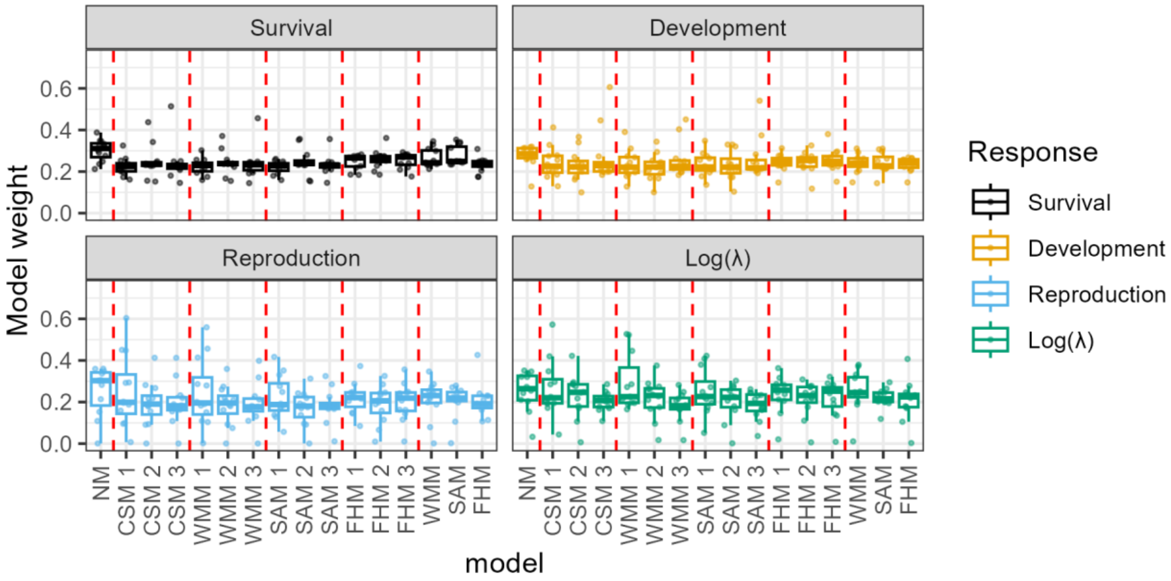

Figure S20. 36-month climate models do not provide predictive advantages when using precipitation data as a predictor. Box and whisker plots and point clouds showing the arcsine transformed model weights on the 12 species with at least 20 years of data, across five model types, three periods (1: year *t*, 2: year *t*-1, 3: year *t*-2), and four response variables. The middle line of the boxplots shows the median, the upper and lower hinges delimit the first and third quartile, and the whiskers extend 1.5 times beyond the first and third quartile. Each point refers to a single model fit.

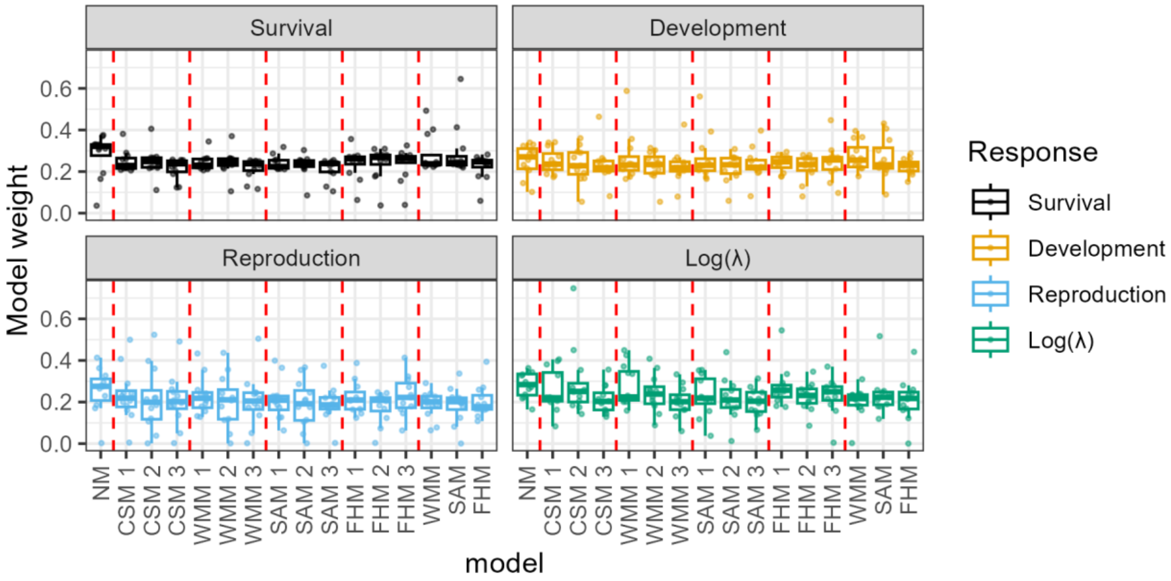

Figure S21. 36-month climate models do not provide predictive advantages when using temperature data as a predictor. Box and whisker plots and point clouds showing the arcsine transformed model weights on the 12 species with at least 20 years of data, across five model types, three periods (1: year *t*, 2: year *t*-1, 3: year *t*-2), and four response variables. The middle line of the boxplots shows the median, the upper and lower hinges delimit the first and third quartile, and the whiskers extend 1.5 times beyond the first and third quartile. Each point refers to a single model fit. The last three columns of each panel refer to models fit on 36 months of data, denoted by the acronyms WMM, SAM, and FHM.

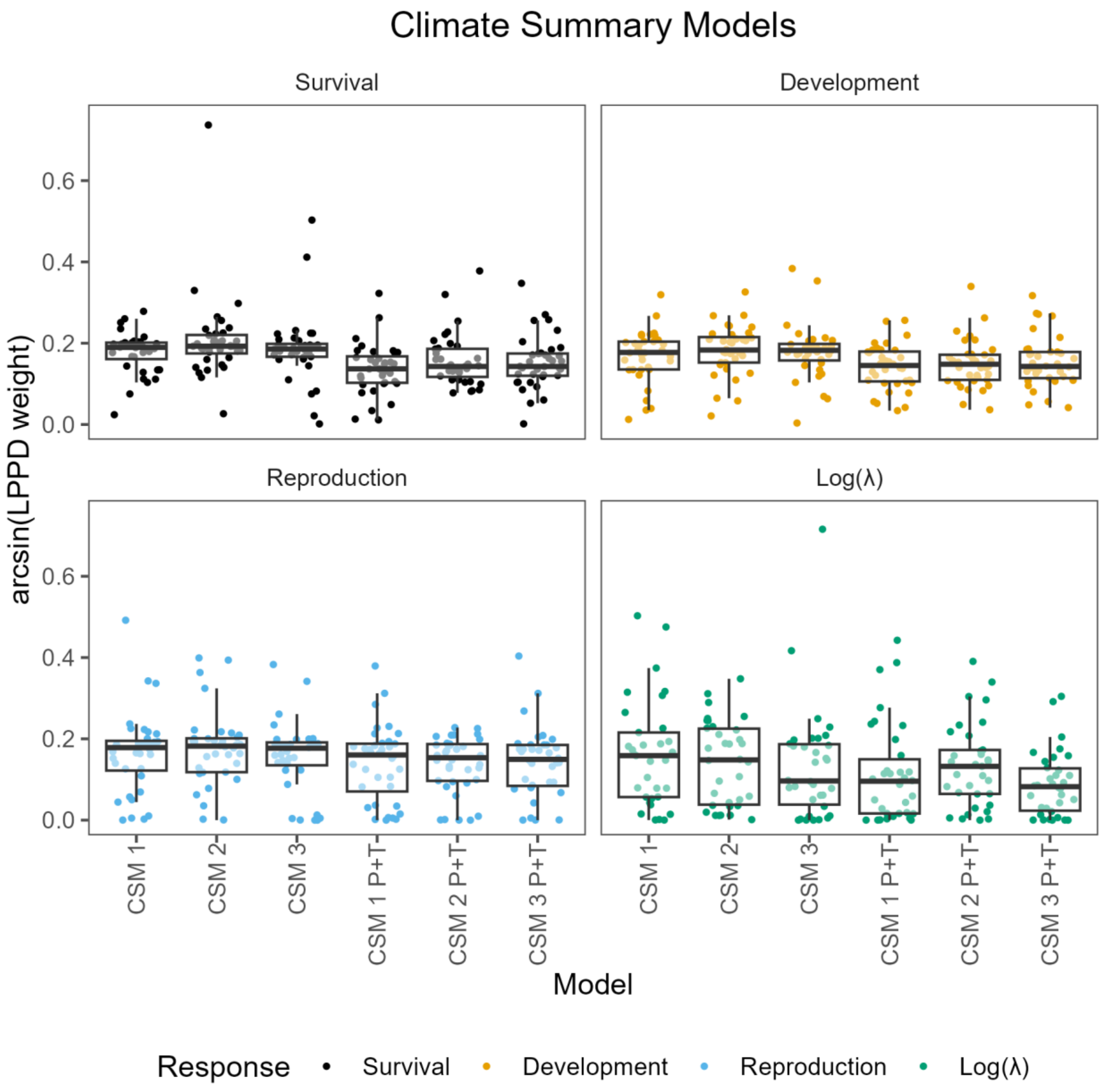

Figure S22. Precipitation plus temperature CSMs do not provide predictive advantages when compared to precipitation-only models. Box and whisker plots and point clouds showing the arcsine transformed model weights for our 34 species, comparing models fit using precipitation as predictors (first three boxplots from the left), to models fit using both precipitation and temperature as predictors (last three boxplots from the right). Each panel refers to a response variable. The middle line of the boxplots shows the median, the upper and lower hinges delimit the first and third quartile, and the whiskers extend 1.5 times beyond the first and third quartile. Each point refers to a single model fit

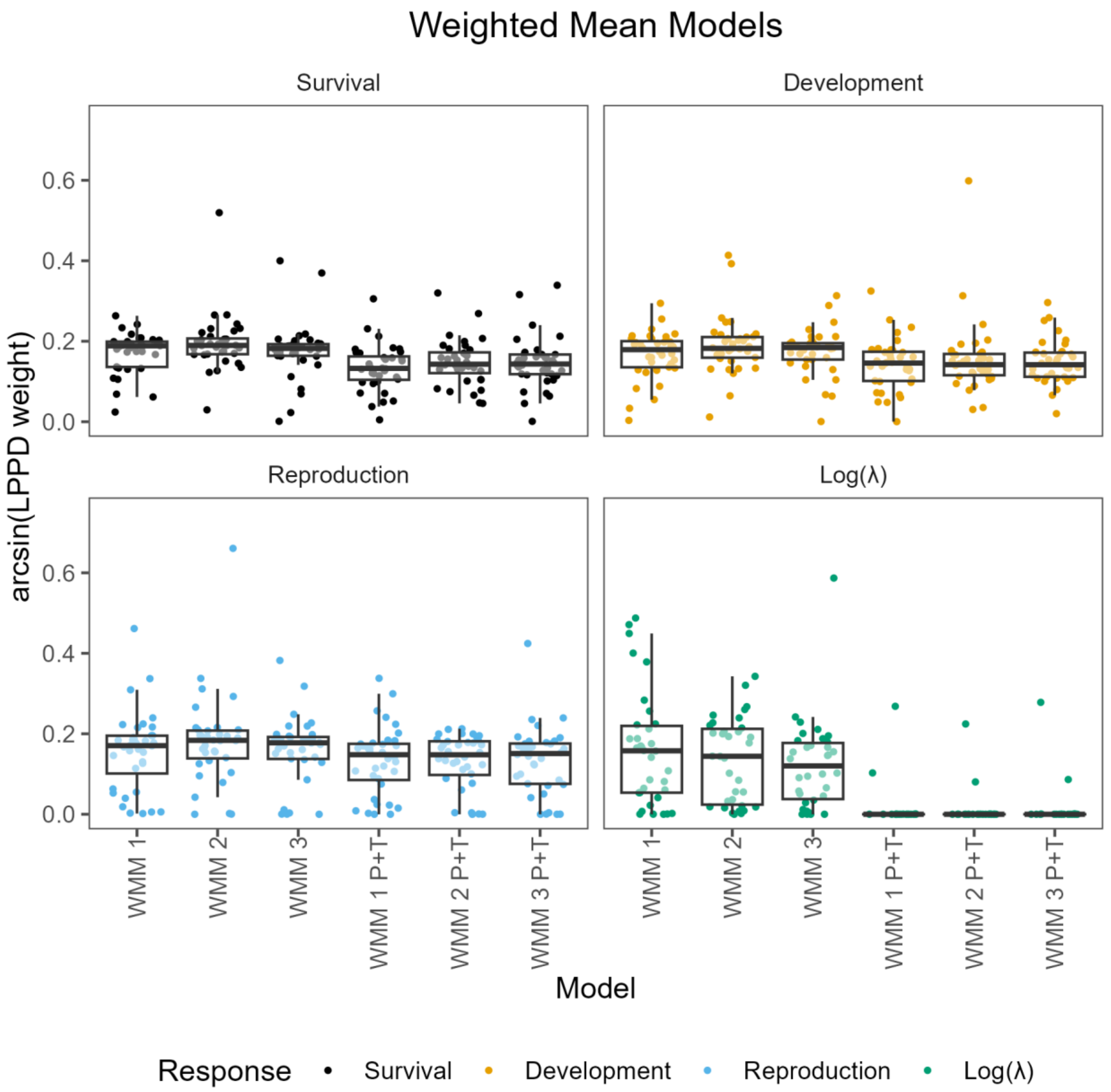

Figure S23. Precipitation plus temperature WMMs do not provide predictive advantages when compared to precipitation-only models. Box and whisker plots and point clouds showing the arcsine transformed model weights for our 34 species, comparing models fit using precipitation as predictors (first three boxplots from the left), to models fit using both precipitation and temperature as predictors (last three boxplots from the right). Each panel refers to a response variable. The middle line of the boxplots shows the median, the upper and lower hinges delimit the first and third quartile, and the whiskers extend 1.5 times beyond the first and third quartile. Each point refers to a single model fit

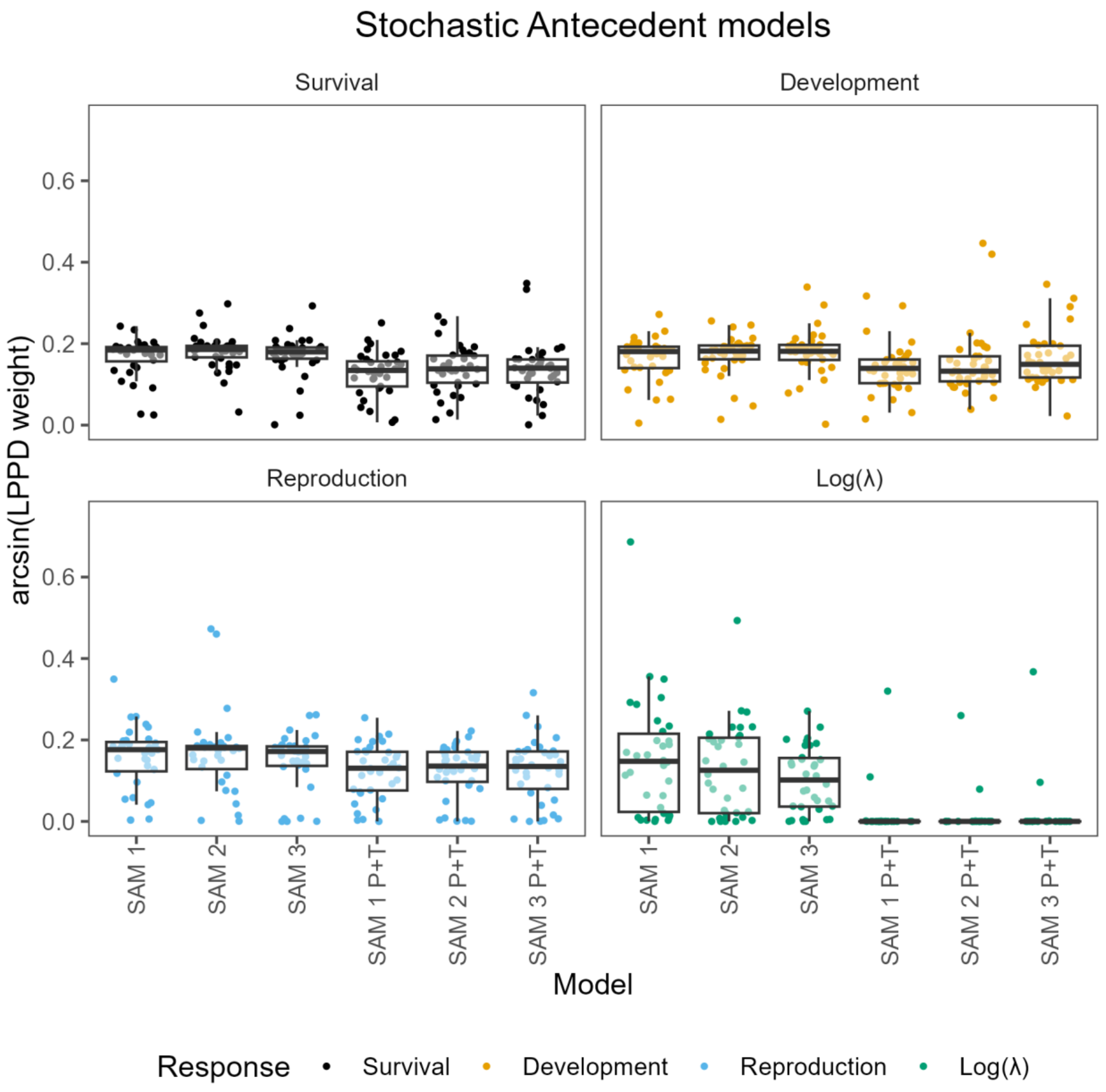

Figure S24. Precipitation plus temperature SAMs do not provide predictive advantages when compared to precipitation-only models. Box and whisker plots and point clouds showing the arcsine transformed model weights for our 34 species, comparing models fit using precipitation as predictors (first three boxplots from the left), to models fit using both precipitation and temperature as predictors (last three boxplots from the right). Each panel refers to a response variable. The middle line of the boxplots shows the median, the upper and lower hinges delimit the first and third quartile, and the whiskers extend 1.5 times beyond the first and third quartile. Each point refers to a single model fit.

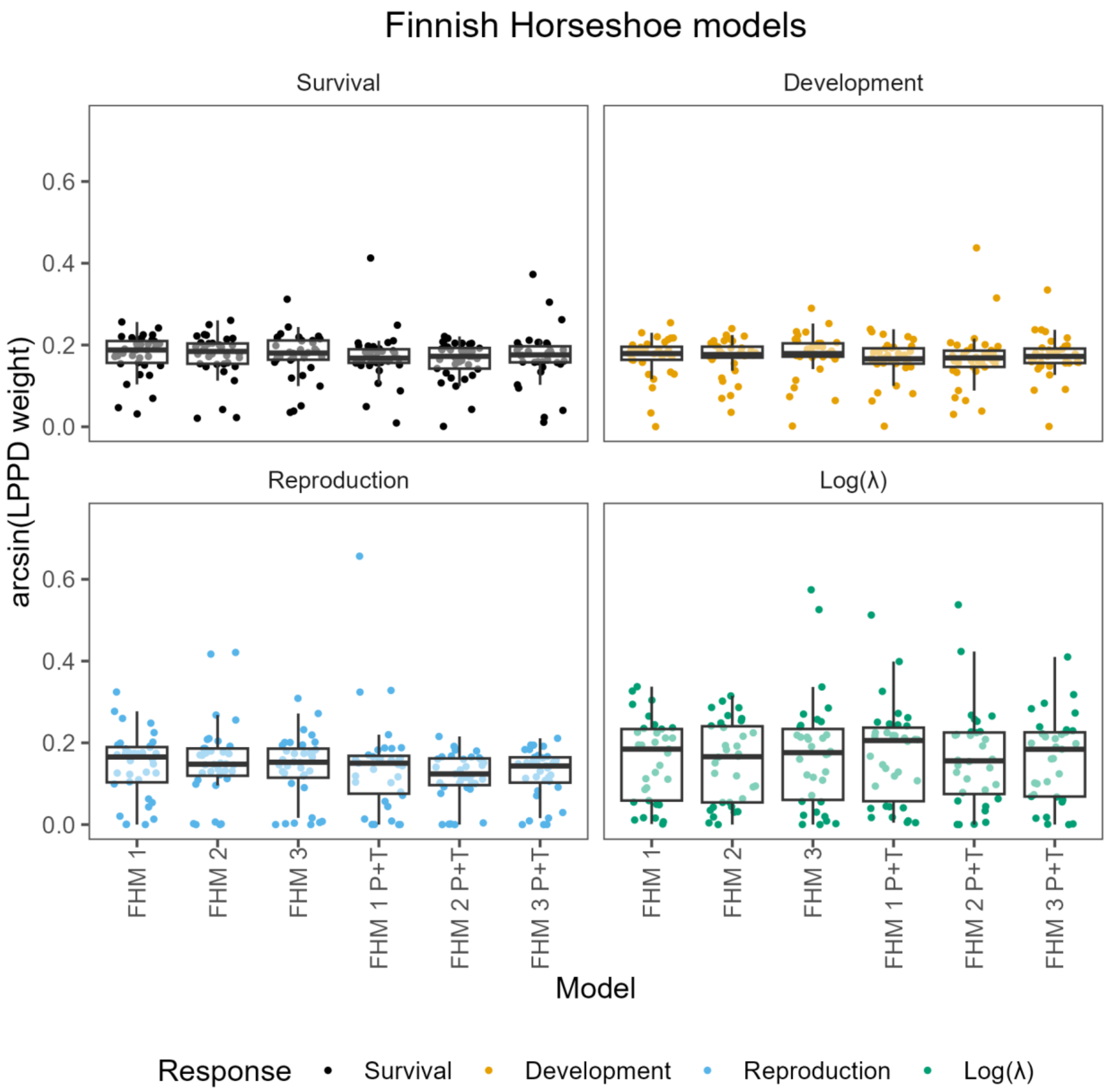

Figure S25. Precipitation plus temperature FHMs do not provide predictive advantages when compared to precipitation-only models. Box and whisker plots and point clouds showing the arcsine transformed model weights for our 34 species, comparing models fit using precipitation as predictors (first three boxplots from the left), to models fit using both precipitation and temperature as predictors (last three boxplots from the right). Each panel refers to a response variable. The middle line of the boxplots shows the median, the upper and lower hinges delimit the first and third quartile, and the whiskers extend 1.5 times beyond the first and third quartile. Each point refers to a single model fit.

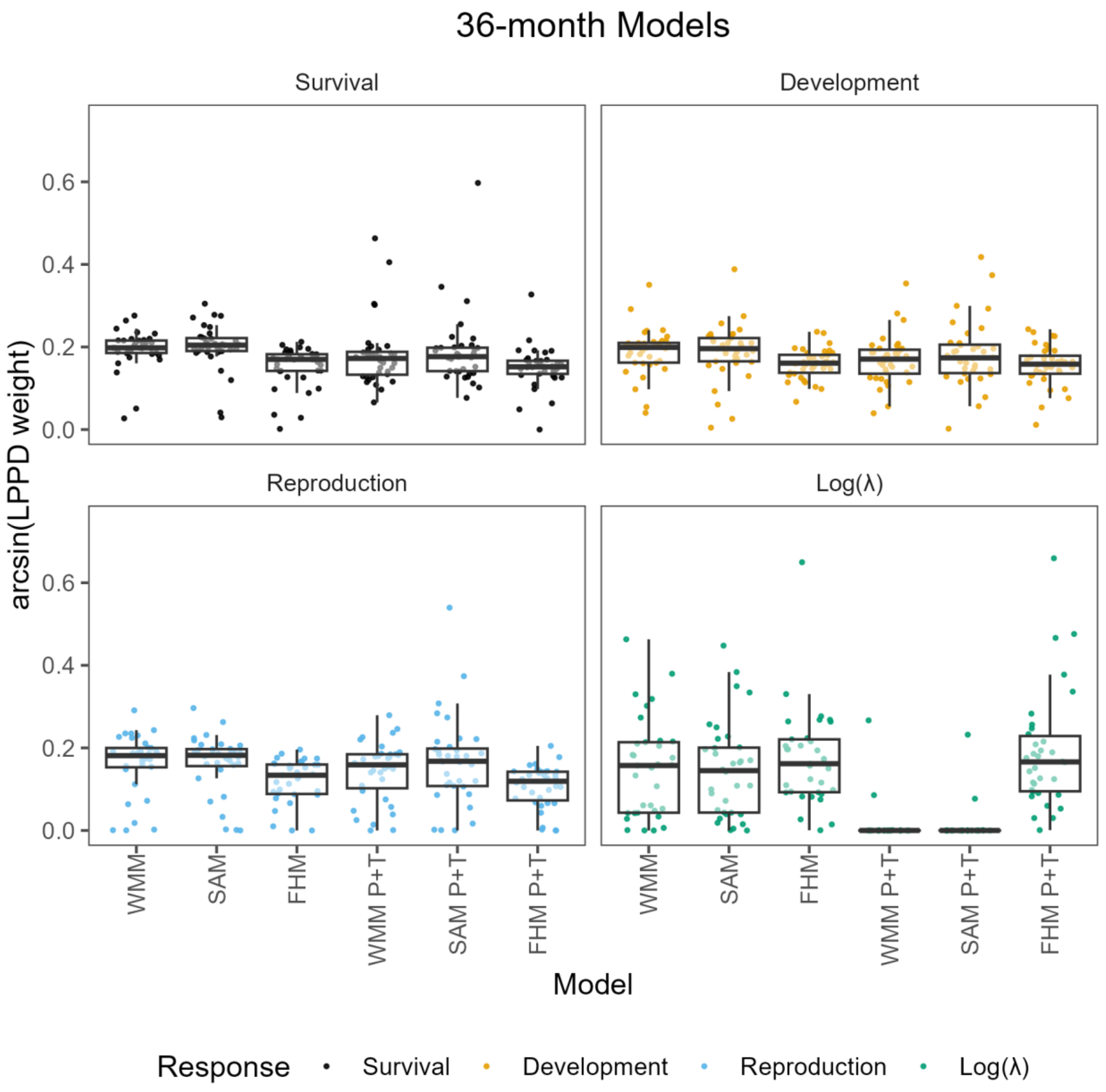

Figure S26. Precipitation plus temperature 36-month antecedent models do not provide predictive advantages when compared to precipitation-only models. Box and whisker plots and point clouds showing the arcsine transformed model weights for our 34 species, comparing models fit using precipitation as predictors (first three boxplots from the left), to models fit using both precipitation and temperature as predictors (last three boxplots from the right). Each panel refers to a response variable. The middle line of the boxplots shows the median, the upper and lower hinges delimit the first and third quartile, and the whiskers extend 1.5 times beyond the first and third quartile. Each point refers to a single model fit.
